# Temporal Proteomics Links the APP London Mutation to Early Cholesterol Remodeling Cell Programs in Human Neurons

**DOI:** 10.64898/2026.10.20.760505

**Authors:** Jesús Muñoz-Estrada, Yuming Jiang, Jesse G. Meyer

## Abstract

Familial Alzheimer’s disease (AD) mutations in amyloid precursor protein (APP) alter amyloid-β (Aβ) secretion profiles, but their effects on neuronal protein and lipid homeostasis remain incompletely understood. To identify early, cell-autonomous phenotypes associated with AD, we used isogenic KOLF2.1J iPSC-derived NGN2 neurons (iNeurons) carrying a heterozygous or homozygous APP-London mutation (V717I) and conducted a longitudinal proteomic study. Results revealed that maturation was the dominant source of proteomic variation, while genotype-dependent effects were pronounced in homozygous APP-London iNeurons. At four weeks, homozygous APP-London iNeurons showed increased APP abundance by proteomics and immunoblot, and increased α-CTF signal by immunoblotting. However, α-CTF/FL-APP and β-CTF/FL-APP ratios from immunoblot did not differ significantly from revertant controls, leaving preferential CTF accumulation unresolved and instead indicating increased steady-state abundance of APP-derived species. Proteomics remodeling in APP-London mutants converged on synaptic, endosomal, and calcium signaling pathways. Notably, homozygous APP-London iNeurons showed a distinct accumulation of cholesterol-biosynthetic enzymes and altered abundance of cholesterol transporters, an increased free/total cellular cholesterol without corresponding induction of SREBP2 target transcripts Pharmacological perturbation of cholesterol metabolism altered Aβ40 and Aβ42 secretion in a compound-specific manner and produced minimal changes in bulk cellular cholesterol, indicating that Aβ secretion was not simply determined by total cholesterol abundance. Finally, we identified SYT4 as a potential γ-secretase target or associated factor that warrants further validation. These findings identify cholesterol dyshomeostasis as an early metabolic phenotype of iNeurons carrying the APP-London mutation, extending the molecular phenotype beyond altered Aβ secretion.

## INTRODUCTION

Alzheimer’s disease (AD) is one of the most common neurodegenerative disorders, and it is estimated that nearly 60 million people aged 65 and older live with clinical AD [1]. Current models suggest that AD progresses through the accumulation of amyloid-β (Aβ) peptides and Tau protein, synaptic dysfunction, and chronic neuroinflammatory responses [2]. These pathological events trigger neuronal death in the brain, clinically characterized by progressive memory loss that affects daily life. Although significant progress has been made in identifying genetic and molecular contributors to AD, the precise relationship of the earliest cellular events that lead to neuronal dysfunction remains complex and is under active investigation. Clinical needs in AD have not been met despite strenuous drug discovery efforts. Currently, a limited number of FDA-approved disease-modifying treatments are effective in patients diagnosed with mild cognitive impairment due to AD.

The advent of human-induced pluripotent stem cell (iPSC) technology has enabled the development of cell models that recapitulate phenotypes observed in patients with AD, while informing relevant biology to elucidate the mechanisms underlying AD pathophysiology [4–6]. iPSC-derived models have demonstrated the utility of this platform for preclinical AD biomarker and drug discovery research [3,3–5]. Importantly, optimizing protocols to induce expression of the transcription factor *NEUROGENIN-2* (*NGN2*) has enabled efficient neuronal conversion of iPSCs, yielding neuron-enriched cultures with cortical excitatory features to investigate early neuronal events that precede AD pathology [6].

In addition to the AD hypothesis driven by amyloid-β (Aβ) peptide imbalance, epidemiological and preclinical studies have reported disturbances in cholesterol homeostasis implicated in AD physiopathology [7]. Brain cholesterol metabolism is linked to both amyloid-β (Aβ) secretion and Tau phosphorylation [8–11]. However, the precise molecular mechanism connecting them remains controversial. For instance, transcriptomic studies in brain regions vulnerable to AD show reduced expression of enzymes involved in de novo cholesterol biosynthesis, consistent with a reduced nuclear translocation of sterol regulatory element-binding protein 2 (SREBP2), a step required to activate gene expression that governs cholesterol biosynthesis [12,13]. In contrast, enzymatic and mass spectrometry-based assays have detected higher levels of both free and esterified cholesterol in AD brain tissue [14,15]. Interestingly, a study in iPSC-derived neurons from AD patients with duplications in the amyloid-β precursor protein gene (APP) found that abnormal accumulation of cholesterol esters enhances Aβ secretion and Tau phosphorylation via independent pathways [16]. This evidence suggests an intricate role of brain cholesterol in the pathophysiology of AD.

In this study, we leveraged an isogenic iPSC-derived neuronal model harboring the APP-London Mutation (V717I), associated with familial Alzheimer’s Disease, to investigate early disease-associated molecular changes in this pathology. We conducted longitudinal proteomic analyses in a set of highly homogeneous NGN2-induced neurons (iNeurons) carrying the APP-London mutation. The set contained homozygous and heterozygous genotypes for this mutation, and as a rigorous control, we used a revertant control cell line, in which the pathogenic allele was corrected to the wild-type allele [17]. In the proteomic dataset, APP-London homozygous iNeurons show more pronounced molecular remodeling over time. In cells with this genotype, we identified a marked increase in the abundance of enzymes involved in cholesterol biosynthesis, as well as cholesterol transporters ABCG1, APOE, LRP1, and LDLR. Consistently, bioluminescence assays indicate elevated free/total cholesterol levels in these cultures, whereas cholesterol esters were undetectable across all differentiated cultures. These changes in cholesterol homeostasis accompanied increased APP abundance at week 4. Pharmacological perturbations of cholesterol metabolism and APP processing altered Aβ secretion while producing minimal changes in bulk cellular cholesterol. Together, these findings characterize APP-London-associated molecular changes in human neurons and support further investigation of the relationship between APP abundance, cholesterol homeostasis, and amyloidogenic processing.

## RESULTS

In this study, we aimed to investigate early molecular events associated with Alzheimer’s disease (AD) by leveraging an isogenic iPSC-derived neuronal cellular platform combined with proteomics. We performed a longitudinal proteomic analysis on NGN2-induced neuronal cultures (iNeurons) carrying the Amyloid Precursor Protein (APP)-London mutation (V717I) (**Fig. 1A**). APP-London iPSC lines were gene-edited using CRISPR-Cas technology using the reference wild-type iPSC line KOLF2.1J, and they were developed by the iPSC Neurodegenerative Disease Initiative (iNDI) [18]. Genotypes of KOLF2.1J-derived isogenic iPSC lines consist of a revertant control line (*APP*^REV/WT^) and APP-specific single-nucleotide variants (*APP*^WT/V717I^) and (*APP*^V717I/V717I^). Of note, the revertant cell line is a strict control in which the disease-causing mutation was reverted to the wild-type allele using the same gene-editing technology. APP-London iNeurons with the three distinct genotypes *APP*^REV/WT^,*APP*^WT/V717I^, *APP*^V7171I/V717I^ are referred to in the manuscript as APP-rev, APP-het, and APP-hom. NGN2 stable integration in KOLF2.1J-derived isogenic iPSC lines was achieved using TALEN-mediated technology as previously described [19]. PCR genotyping and karyotype analysis of NGN2 KOLF2.1J-derived isogenic single-cell clones across APP-rev, -het, and -hom genotypes indicated monoallelic integration of NGN2 into the AVVS1 locus and the absence of genetic abnormalities (**Fig. S1A, B**). We next generated iNeurons using an optimized protocol that yields negligible numbers of progenitors or glial cells, resulting in neuronal-enriched cultures [20]. This experimental strategy ensures that the observed proteomic signals primarily reflect neuronal cells and enables the identification of neuronal-specific phenotypes associated with the APP-London mutation.

**Figure 1.**
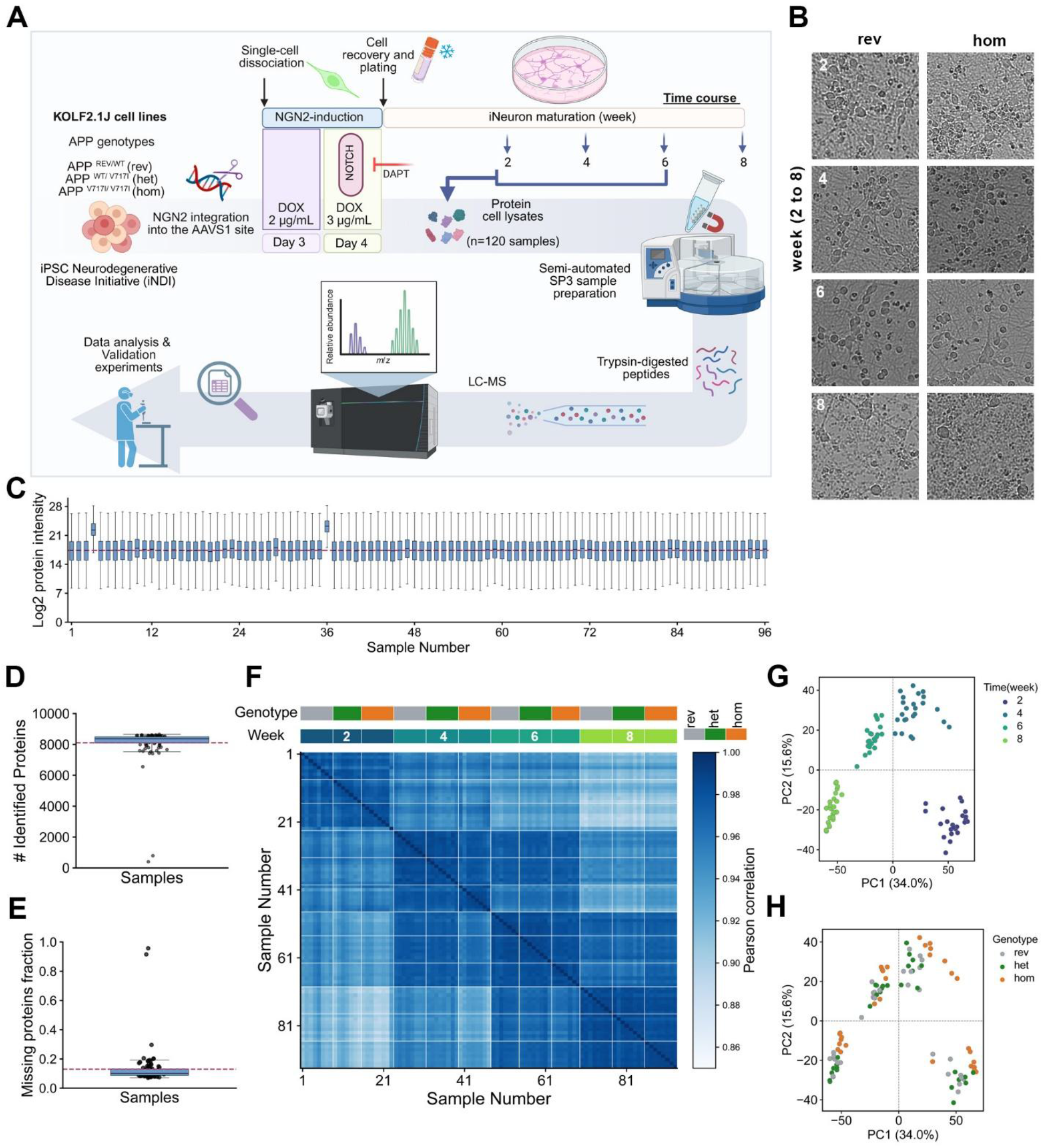
Longitudinal proteomic profiling of an isogenic iNeuron set harboring the APP-London mutation (V717I). **(A)** Schematic representation of the experimental design in CRISPR-Cas gene-edited isogenic KOLF2.1J iPSC lines harboring *APP*^REV/REV^, *APP*^WT/V717I^, and *APP*^V717I/V717I^, designated as APP-rev, APP-het, and APP-hom. APP-rev is a strict control in which CRISPR-Cas technology reverted the pathogenic allele to the wild-type allele. We performed TALEN-mediated stable integration of NGN2 at the AAVS1 locus in the three iPSC lines, followed by a 4-day period of NGN2 induction and neuronal maturation of iNeurons across weeks 2, 4, 6, and 8. By week 8, iNeurons were lysed and semiautomated proteomic sample preparation was performed using the SP3 magnetic-bead method on a KingFisher Flex platform for LC-MS analysis. We then performed data analysis and validation experiments. **(B)** Representative brightfield microscopy images of iNeurons at weeks 2, 4, 6, and 8 post-NGN2 induction. At week 4, neuronal morphologies are evident, whereas at week 8, cell somata are clearly reduced (neuronal loss). **(C)** Boxplot of Log2-transformed protein intensities across all 96 samples (n=4 time points, n=8 technical replicates, n=3 genotypes) showing baseline uniformity (mean ± SD: 17.47 ± 0.8), mean is indicated with the burgundy horizontal dashed line. **(D)** Boxplot showing the total number of unique identified proteins per sample, highlighting a depth of ∼8,000 proteins across samples and two clear outliers with a protein count below 1,000 (excluded from downstream analysis). **(E)** Fraction of missing values across the proteomic dataset (∼13%), demonstrating robust data depth and sample reproducibility. **(F)** Heatmap of sample-to-sample Pearson correlation coefficients across the three APP genotypes and four time points, clearly driven by the temporal stage of neuronal maturation rather than APP allele status. **(G)** Principal Component Analysis (PCA) screening maturation time points and **(H)** APP genotypes of iNeurons.

Semiautomated proteomic sample preparation of iNeurons maintained for 2, 4, 6, and 8 weeks after NGN2 induction was performed using the SP3 method and a KingFisher Flex Magnetic Particle Processor (**Fig. 1A**) [21]. **Figure 1B** shows representative brightfield images of these cultures, in which neuronal morphologies are observed, particularly in weeks 4 and 6. At week 8, a reduced number of cell somas is also observed in the field of view, suggesting neuronal loss in long-term cultures. Analyses of the derived iNeuron proteomic dataset (**Supplementary Tables 1, 2, and 3**) across the three London-APP genotypes (rev, het, and hom) spanning four time points (weeks 2, 4, 6, and 8) with eight technical replicates per condition showed a consistent distribution of Log2-transformed protein intensities (96 samples total), except for two clear outliers (overall mean ± SD, 17.47 ± 0.8) (**Fig. 1C**, burgundy dashed line). The number of proteins identified across samples was ∼8,106, except for the outliers, in which fewer than 1,000 proteins were identified (**Fig. 1D**). The fraction of missing proteins is modest (∼13%), indicating robust protein identification depth across samples. For downstream analysis, we filtered samples using a 15% deviation threshold from the average number of proteins identified, excluding 3 samples: a replicate from APP-rev-6, -hom-6, and het-2 (genotype-week). These sample metrics indicate high, consistent proteome coverage across the dataset, including the week 8 timepoint, where a reduction in the number of cells is observed. Sample-to-sample Pearson correlation analysis showed high overall concordance across the dataset, with the strongest correlations driven primarily by timepoint across the three APP-London genotypes (**Fig. 1F**), indicating that time is a major driver of proteomic variation in these cultures. To further identify the sources of variance, we performed principal component analysis (PCA). The first two principal components account for almost half of the variance and reveal a clear temporal trajectory, with samples forming separate clusters along PC1 by maturation time point (**Fig. 1G**). In contrast, genotype-dependent separation of the APP-London iNeurons was mildly apparent, particularly for the APP-hom condition, indicating that while genotype contributes to proteomic variation, its effect is secondary in the dataset (**Fig. 1H**).

Proteomic molecular characterization of APP-London iNeurons across time points (2, 4, 6, and 8 weeks) was analyzed by the relative abundance of specific cell markers (**Fig. 2A**). Using z-scores of selected protein markers and a minimum detection threshold of 50% across replicates, we observed a similar temporal molecular pattern in iNeurons with the three distinct APP-London genotypes. Pluripotency markers SOX2 and OCT4 were absent from the dataset, and PODXL was detected only at week 2, indicating residual expression of these markers in progenitor cells transitioning to post-mitotic neurons [20]. Conversely, markers of mature neurons, including RBFOX3 (NeuN), NEFM, SNAP25, and SYN1, showed a high relative abundance in week 4. Expression of markers indicating cortical excitatory identity of iNeurons (i.e., GLS, SLC1A6, CAMK2A) was upregulated during weeks 2 and 4. Glutamatergic identity is supported by our previous immunofluorescence analyses in KOLF2.1J-derived iNeurons [20]. Expression of cholinergic neuronal markers was high during weeks 2 and 4, consistent with independent proteomic data of KOLF2.1J-derived iNeurons [22], suggesting a partial co-activation of cholinergic programs mediated by NGN2 [23]. Unexpectedly, the relative expression of peripheral neuronal markers was upregulated at week 8, raising the question of whether the spontaneous reduction in cell numbers observed at that time primarily reflects the loss of neurons with glutamatergic identity or the activation of peripheral neuronal programs in long-term cultures (**Fig. 1B**).

**Figure 2.**
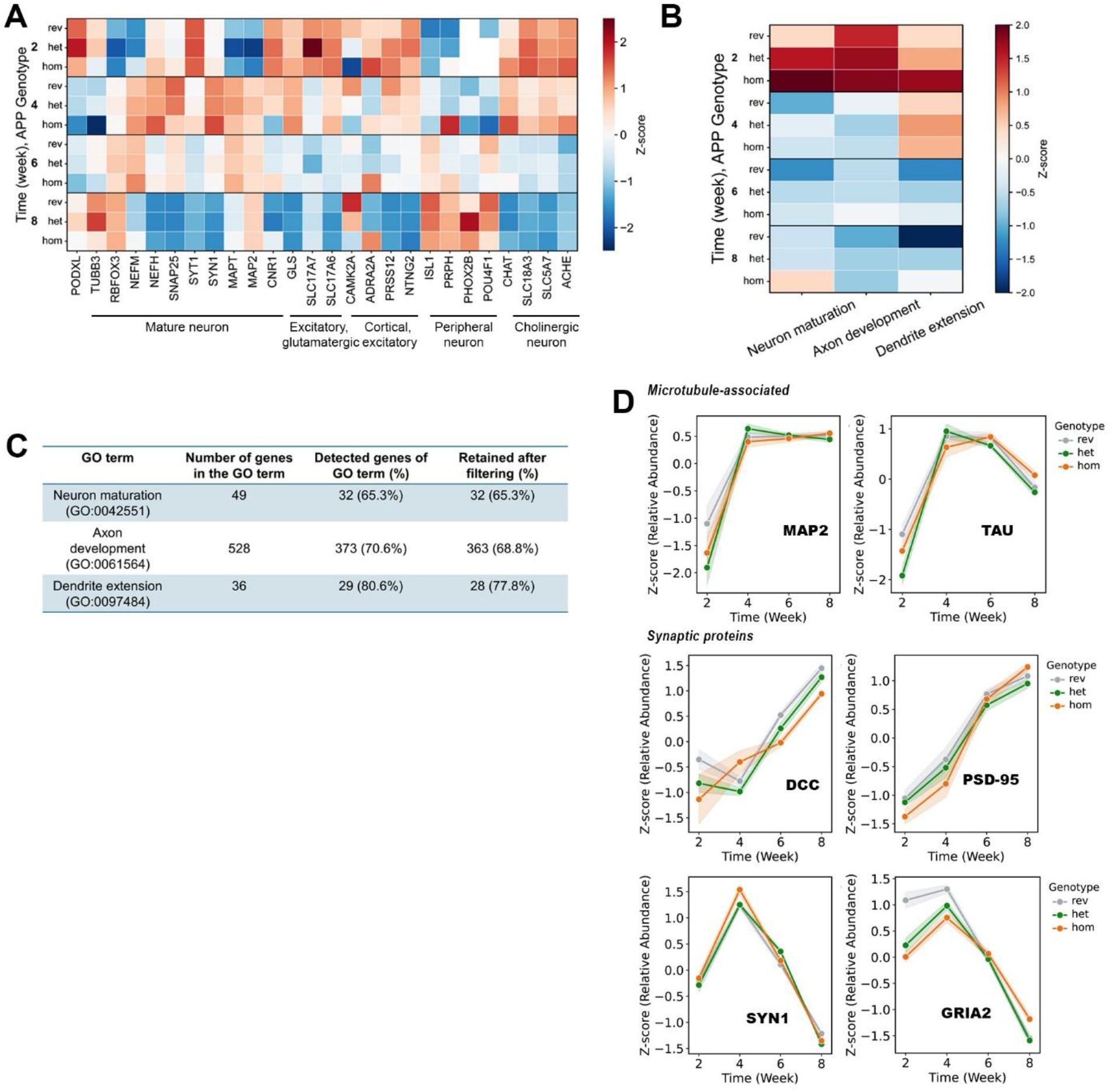
Molecular Characterization of iNeurons carrying the APP-London mutation (V717I). **(A)** Heatmap of protein abundances (z-scores) of canonical lineage markers across the 2-, 4-, 6-, and 8-week time courses, including pluripotency markers, mature neuronal markers, glutamatergic and cortical neuronal markers, peripheral neuronal markers, and cholinergic program markers for each APP genotype: APP-rev, APP-het, and APP-hom. **(B)** Heatmap of relevant neuronal Gene Ontology (GO) terms, including Neuron Maturation, Axon Development, and Dendrite Extension at each time point across the 8-week time course for the three genotypes. **(C)** Table with proteome coverage and filtration metrics for the targeted GO terms in (B). Of note, the analysis was performed only on proteins quantified in at least 6 of the 8 technical replicates (≥75%) within each experimental group. **(D)** Protein expression z-scores (y-axis) at each time point across the 8 weeks (x-axis) for canonical neuronal maturation markers (MAP2, TAU) and synaptic proteins (DCC, PSD-95, SYN1, GRIA2) for each APP genotype. Points at each time point represent the mean Z-score; the shaded region in all trajectory plots indicates the standard error of the mean (SEM).

To assess the dynamics of neuronal-specific biological processes iNeurons, z-scores were calculated for three Gene Ontology (GO) terms across time: ‘Neuron Maturation’, ‘Axon Development’, and ‘Dendrite Extension’. We observed that the three biological programs were upregulated in week 2. Interestingly, the ‘Dendrite Extension’ program sustained the high protein expression through week 4, whereas scores for all three pathways decreased by Weeks 6 and 8 (**Fig. 2B**). **Figure 2C** summarizes the coverage of these GO terms in the iNeurons proteomic dataset, retaining only proteins detected in at least 6 out of 8 technical replicates within each experimental group (75%). Overall, this analysis suggests that molecular programs driving the specialization of these iNeurons, regardless of APP genotype, are established by week 4 of differentiation, consistent with previous timelines for establishing NGN2-induced functional neurons [4,22]. We then analyzed the expression of canonical neuronal maturation markers over the 8-week differentiation period using z-score-standardized abundance. As expected, the microtubule-associated proteins MAP2 and TAU show a steep increase in relative expression from week 2 to week 4, followed by a plateau (**Fig. 2D**). Interestingly, DCC and PSD-95, which are enriched in the postsynaptic density (PSD) and participate in glutamatergic neurotransmission in forebrain neurons, show a gradual increase over the 8-week differentiation period, suggesting a dynamic structural synaptic remodeling in iNeurons maintained for an 8-week period [24,25] (**Fig. 2D**). Although, inference of synaptic activity in iNeurons should be interpreted with caution, as other synaptic proteins are not detected (i.e. NTN1 ligand of DCC) or peak their expression at week 4 (SYN1, GRIA2) (**Fig. 2D)**.

We then examined the APP relative expression profile in the iNeuron proteomic dataset, and results showed the highest expression at week 2 in the three APP-London genotypes, implicating its participation in early development during neuronal differentiation and axonal outgrowth (**Fig. 3A**). A lower expression beyond week 2 can be associated with the establishment of functional excitatory synapses (**Fig. 3A**) [26–28]. Despite the trend toward reduced APP levels consistent with maturation of iNeurons, APP-hom cells show a clear increase in abundance at week 4 relative to both APP-rev and APP-het genotypes (**Fig. 3A**, red arrowhead). Quantification of Log 2-centered abundances (average baseline=0) that reflect the variance of the raw dataset reveals a robust accumulation of APP holoprotein in APP-hom iNeurons at week 4 (**Fig. 3B**, **P-adj <0.01) compared to either APP-het or -hom. Interestingly, we also detected a marked accumulation at week 4 of the two APP orthologues, APLP1 and APLP2, in APP-hom iNeurons (**Fig. S2**), supporting that the APP gene family follows similar expression patterns [29].

**Figure 3.**
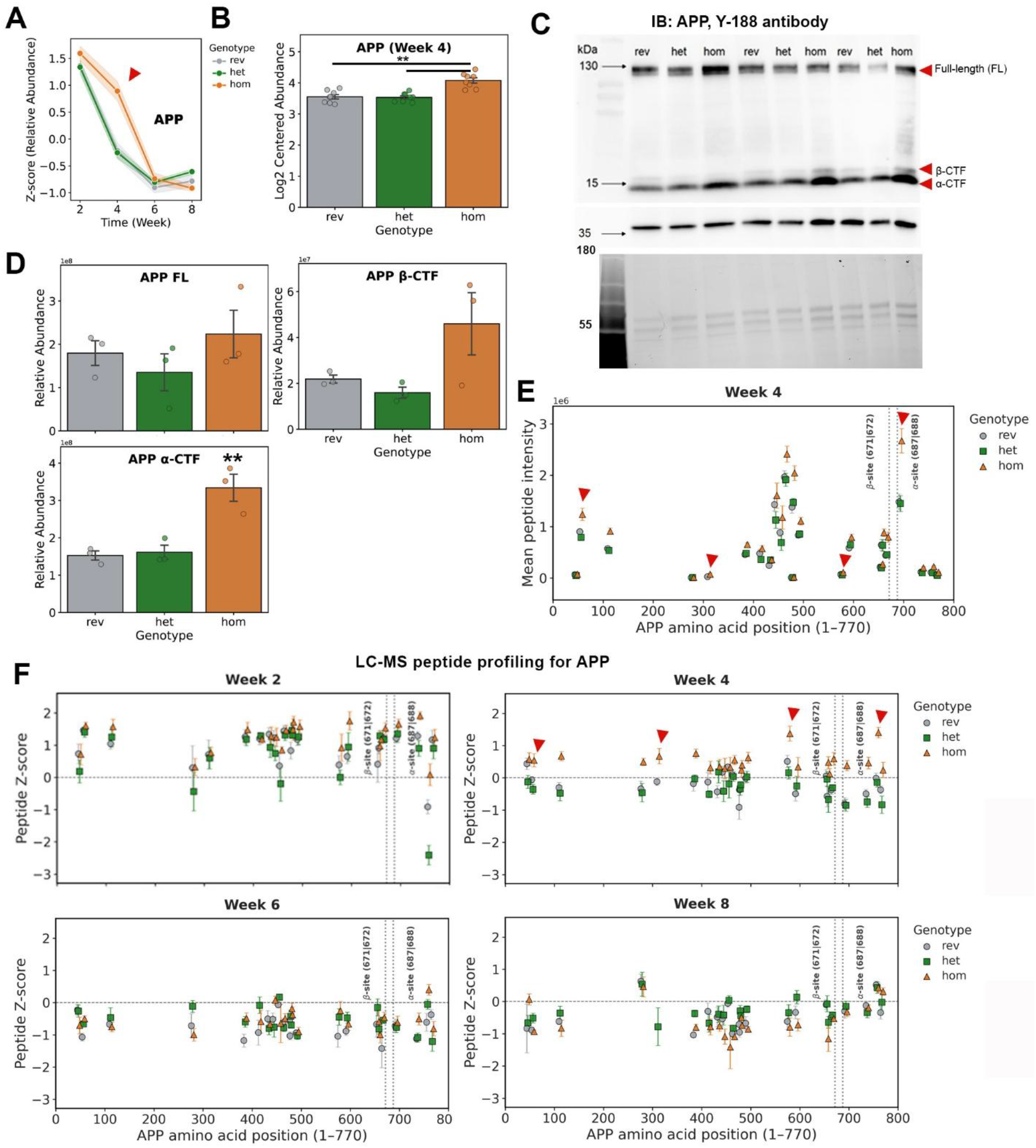
APP expression analysis in iNeurons carrying the APP-London mutation. **(A)** Protein expression z-scores and **(B)** log2-centered relative abundance profiles of full-length APP across the 8-week differentiation period and at week 4 for each APP genotype. In (A), the red arrow indicates a significant accumulation of APP expression distinct to the APP-hom genotype at week 4 (red arrow), and in (B), **P < 0.01. **(C)** Western blot analysis of iNeuron whole lysates at week 4 from the three isogenic lines, probed with an anti-APP (Y188) antibody. The band molecular weights correspond to full-length APP (FL-APP) and cleaved β-CTF and α-CTF fragments. The image also showed the GAPDH loading control and the stain-free gel used for normalization. **(D)** Densitometric analysis of Western blot signals (APP-FL, α-CTF, and β-CTF) for the three genotypes; APP-rev, APP-het, and APP-hom (n=3 independent biological replicates; **P < 0.01). The shaded region in all trajectory plots indicates the standard error of the mean (SEM), and error bars in bar plots indicate the standard error of the mean (SEM). **(E)** LC-MS peptide intensity profiling at week 4 across the full-length human APP sequence (1-770 aa) at the indicated time points. Each symbol represents the mean intensity. Error bars indicate the standard error of the mean (SEM). Each dot denotes an individual peptide plotted at its central sequence coordinate (amino acid midpoint). **(F)** Peptide-specific Z-scores across the indicated time points; each symbol represents the mean peptide Z-score across biological replicates, with error bars indicating SEM. In both panels (E and F), dotted vertical lines mark the canonical BACE1 cleavage site (β-site: Met671|Asp672) and α-secretase cleavage site (α-site: Lys687|Leu688).

Western blot analysis of week-4 iNeuron lysates using the Y188 antibody detected full-length APP (FL-APP) and two lower-molecular-weight bands consistent with APP C-terminal fragments (CTFs) (**Fig. 3C**) [30]. After normalization to total lane protein, APP-hom cultures showed higher mean signals for FL-APP and β-CTF, with nominal evidence for increased α-CTF relative to APP-rev (**Fig. 3D**, nominal P < 0.05, n = 3). However, α-CTF/FL-APP and β-CTF/FL-APP ratios calculated within individual samples did not differ significantly from APP-rev (**Fig. S2C**). Peptide-level LC-MS analysis mapped to the canonical 770-amino-acid APP sequence showed higher abundances for several peptides distributed across the sequence in APP-hom cultures at week 4 compared with APP-rev and APP-het (**Fig. 3E**, red arrowheads). Peptide-specific z-scores made these differences more apparent (**Fig. 3F**, week 4), whereas peptide profiles were generally more similar across genotypes at weeks 2, 6, and 8. Because C-terminal peptides are shared by FL-APP and its cleavage products, these measurements do not independently resolve CTF abundance. Together, the results support increased APP abundance in APP-hom cultures at week 4 but do not establish preferential accumulation of either CTF or a selective shift in α-or β-secretase processing. The increase in APP abundance is consistent with previous reports of elevated FL-APP in iPSC-derived neurons from patients carrying the APP-London mutation (V717I) [31].

We then analyzed the expression proteins belonging to the ADAM (a disintegrin and metalloprotease) family (ADAM9, ADAM10, ADAM16, ADAM17) and β-secretase (BACE1) enzymes that participate in the generation of APP CTFs and observed similar z-scored expression trends across the three APP-London genotypes over time (**Fig. 4A-B**). The highest expression of the main proteins with α-secretase activity, ADAM9 and ADAM10, in iNeurons is observed at week 2, followed by a decrease at week 4 and a gradual increase in ADAM10 expression at subsequent time points. Targeted analysis of Log2-centered abundance of individual proteins restricted to week 4 showed no significant changes for ADAM10 and ADAM9 across the three genotypes (**Fig. S2A**). The lowest expression of ADAM16 and ADAM17 is observed at week 4 and follows the ADAM10 expression trend across maturation time points in iNeurons [32]. Although α-secretase activity for APP has been detected in ADAM9, ADAM10, and ADAM17, ADAM10 is the major α-secretase for APP [32,33]. ADAM10 has the highest expression at early time points, which may explain its role in neurite outgrowth during development, whereas at later stages of neuronal maturation, it may be involved in the structural organization and function of glutamatergic synapses [34–36].

**Figure 4.**
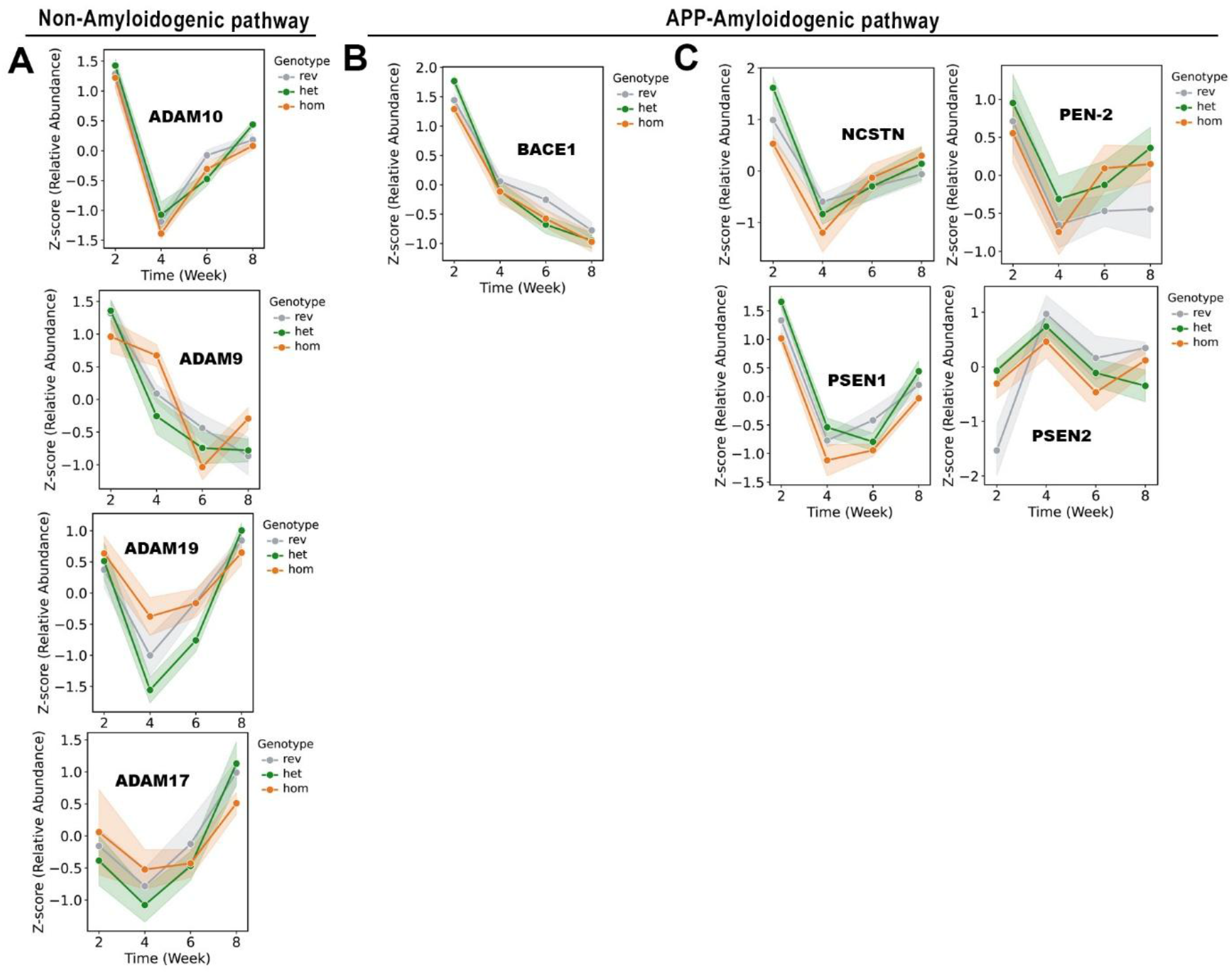
Proteomic remodeling of the amyloidogenic and non-amyloidogenic pathway in APP-London iNeurons across maturation timepoints. **(A)** Protein expression z-score profiles (y-axis) at each time point over 8 weeks (x-axis) for proteins belonging to the ADAM family (ADAM10, ADAM9, ADAM19, ADAM17) in APP-rev, APP-het, and APP-hom iNeurons. Each point represents the mean z-score, and lines connect the mean values over time. **(B)** Protein z-score trajectories for the major β-secretase enzyme BACE1 across the 8-week time course in the three APP genotypes. **(C)** Z-score trajectory plots for members of the γ-secretase complex (NCSTN, PEN-2, PSEN1, PSEN2), showing peak expression levels during early differentiation of iNeurons in the three APP genotypes.

For BACE1, the major β-secretase in vivo, the relative expression across time points follows the APP trend: upregulation at week 2, followed by a steep decrease, which can be explained by its role in neurite outgrowth [37] (**Fig. 4B**). Unnoticeable changes in relative expression were detected for BACE1 across genotypes (**Fig. S2B**). BACE2 was not detected in the dataset, and its activity may not contribute to Aβ pathology in AD [38] (PMID: 19968762). For the γ-secretase components NCSTN, PEN-2, PSEN1, and PSEN2, the highest expression is observed at week-2 (**Fig. 4C**), as is the case for ADAM10, BACE1, and APP. These may suggest active involvement of components of γ-secretase in regulating neuronal specialization, although the interpretation is limited given its multiple subcellular localizations and activity [39].

Our results from the iNeurons and previous studies indicate that the APP-London mutation preferentially alters the APP amyloidogenic pathway. To better understand molecular alterations associated with these biochemical imbalances in APP metabolism, we implemented a two-way factorial linear model with interaction terms (Intensity ∼ Timepoint x Genotype) on the 8-week iNeuron maturation time-course proteomic dataset. **Supplementary Table 4** shows the results. Of the 7,319 screened proteins, statistical results indicate that the Timepoint nodule exerts a main effect on the dataset, followed by a Genotype effect (7023 and 3583 statistically altered proteins according to the statistical cutoffs, respectively; P-adj <0.05) (**Fig. 5A**), consistent with PCA results (**Fig. 1G** and **H**). Ordered heatmap of the top-200 time-regulated proteins by their peak expression timing and temporal trajectory in APP-rev controls reveals similarities in maturation trajectories across APP-London iNeurons with clear divergences in the APP-hom genotype at week-4 (**Fig. 5B**). The (APP-hom vs APP-rev) and (APP-het vs APP-rev) contrast results (3277 and 244 significant proteins, respectively) indicate that the Genotype effects are largely mediated by APP-hom. Differential expression analysis of the pairwise contrasts (P-adj) <0.05), with APP-rev used as the baseline control, revealed that only STOM and PCDHGB4 exhibit a marked accumulation (∼2-fold increase) in both APP-het and APP-hom (**Fig. S3A-B**). APP-het iNeurons display a specific dysregulation in structural proteins (protocadherins, PDCH) (**Fig. S3B**). In contrast, the APP-hom exhibited broader alterations across a diverse group of proteins, including those involved in ion homeostasis and neuronal survival, such as CALB2, S100A13, MRS2, and STOM (**Fig. S3A**). To assess gene-dosage effects across the APP-London iNeurons, we performed a genotype concordance analysis mapping the molecular effect size (log2 FC) of APP-het vs APP-hom against the APP-rev (baseline). Of 7,319 proteins, only 83 exhibited significant temporal dysregulations in APP-het. In stark contrast, out of the 3,277, 3,116 proteins showed dysregulation in the APP-hom condition (**Fig. S3 A-B**), and only 161 proteins displayed directional concordance between mutant genotypes by co-localizing in quadrants I and III (**Fig. 5C**). In addition, only 30 proteins moved in opposite directions, suggesting that a higher number of proteins display directional concordance and reflecting that there is more evident molecular rewiring in APP-hom iNeurons. Further analysis facilitated by the Alzheimer’s Disease Open Targets Platform, which includes GWAS catalog studies, revealed that several directionally concordant proteins have established links to AD-relevant processes, including amyloid and tau pathology and glutamatergic synaptic function. For example, INPPL1/SHIP2 has been implicated in linking Aβ-associated signaling to tau pathology, PLXNA4 has been associated with AD risk and tau phosphorylation, and CAMK2A and GRIA2 participate in glutamatergic synaptic pathways disrupted by oligomeric Aβ (**Fig. 5C**) [40–44]. Functional enrichment analysis of the protein concordance results of APP-het and APP-hom revealed a strong overrepresentation of neuronal-related and glutamatergic synaptic pathways (**Fig. S3C**) [45]. Molecular function analysis indicated enrichment for calmodulin and calcium-binding proteins, consistent with calcium signaling perturbations in Alzheimer’s disease (**Fig. S3C**) [46]. Interestingly, enrichment analysis of the APP-hom revealed temporal regulation of biological pathways strongly associated with AD pathology [16,30,47]. Among these pathways we found endosomal trafficking, synapse organization, mitochondrial respiratory chain, cholesterol biosynthesis, and Alzheimer’s Disease pathways (**Fig. S3D**). A full list of the specific and shared proteins dysregulated in APP London homozygous and heterozygous iNeurons is provided (**Supplementary Table 5**).

**Figure 5.**
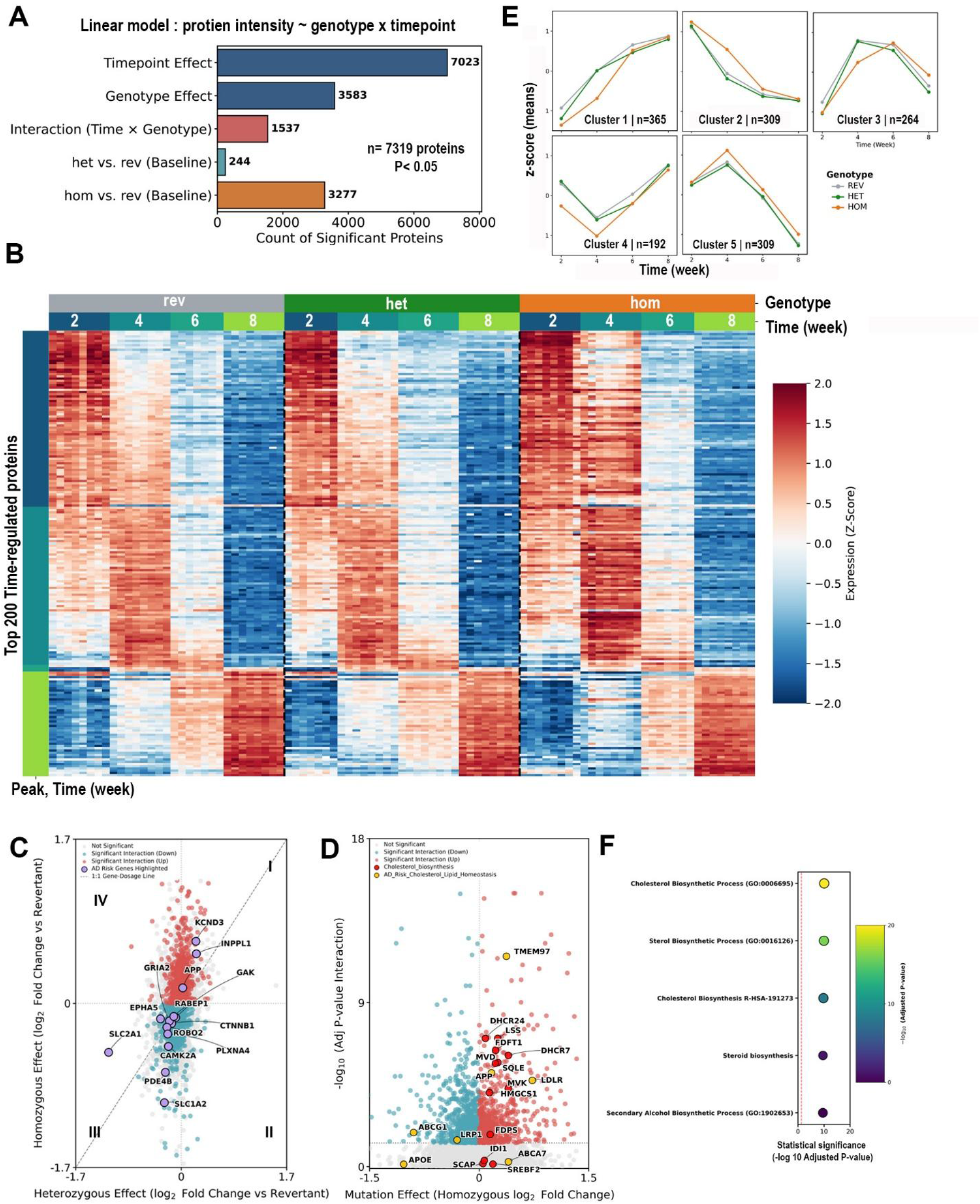
Linear model results of the longitudinal proteomic dataset from iNeurons harboring the APP-London mutation. **(A)** In the chart, horizontal bars indicate the number of significant proteins (P-Adj < 0.05) for each node. Model terms include the main effects of Timepoint and Genotype, the (Timepoint x Genotype) interaction, and Genotype contrasts (APP-het or APP-rev vs APP-rev). **(B)** Peak-ordered temporal expression heatmap of the top 200 time-regulated proteins. Screened and selected proteins with the lowest adjusted P-value for the time-point effect (P-Adj Timepoint). Proteins are arranged along the vertical axis by their chronological peak of maturation (week) and temporal center of mass in the APP-rev control line. Z-scores represent relative, row-standardized protein abundance across genotypes (APP-rev, APP-het, and APP-hom) and maturation time points (Weeks 2, 4, 6, and 8). Warmer colors indicate relatively higher abundance, whereas blue colors indicate relatively lower abundance for each protein. **(C)** Scatter plot of log2 fold changes (log2 FC) for APP-het vs APP-rev (x-axis) against APP-hom vs. APP-rev (y-axis) (log2 FC=0; APP-rev serving as a baseline). Each protein represents a single protein, and the diagonal line indicates equal effect sizes across the two contrasts (APP-het and APP-hom). Proteins with significant timepoint x genotype interaction (P-Adj <0.05) are shown in red (positive log2 FC) or blue (negative log2 FC). Annotated purple dots represent Alzheimer’s Disease risk genes. **(D)** Volcano plot showing the APP-hom genotype effect (APP-hom vs. APP-rev log2 fold change; x-axis) and the significance of interaction (Time x Genotype) (−log10 P-Adj value; y-axis). The vertical dotted line indicates no change relative to the APP-rev group (log2FC = 0), and the horizontal dotted line denotes the significance threshold (P-Adj = 0.05). Significant proteins are shown in red (positive log2FC) or blue (negative log2FC), whereas nonsignificant proteins are shown in gray. Proteins involved in cholesterol biosynthesis are colored red, and proteins involved in cholesterol transport are highlighted in yellow. **(E)** Profile trajectories of K-means clustering (K = 5) tracking the distinct temporal trajectories of 1,439 quantified interaction proteins in APP-London iNeurons. Each panel shows the mean z-score trajectory for proteins within a cluster across experimental weeks 2, 4, 6, and 8, and the number of proteins identified with similar expression profiles across three APP genotypes. Proteins with missing values in any sample were excluded prior to clustering. **(F)** Bubble plot displaying significantly enriched biological terms with proteins assigned to Cluster 2. Top 5 enriched terms (y-axis) with significance (x-axis). Bubble color reflects adjusted P values, with lighter colors indicating greater statistical significance. Terms are ordered by enrichment significance, and the vertical red dashed line denotes the significance threshold corresponding to a P-Adj value of 0.05.

To further analyze the marked phenotypes in APP-hom neurons associated with AD pathology and to evaluate variations in protein abundance alongside significant time-course alterations. We performed a Genotype-by-Time interaction analysis, contrasting the APP-hom effect size (log2 FC, x-axis, n=3,277) with P-adj values (<0.05) from the Interaction (Time x Genotype) term (y-axis, n= 1537) in the linear model (**Fig. 5D**). The analysis reveals that cholesterol biosynthetic enzymes, including HMGCS1, MVK, MVD, FDPS, SQLE, LSS, FDFT1, DHCR7, DHCR24) (**Fig. 5D**, red dots) as well as proteins related to cholesterol homeostasis (LDLR, TMEM97, APP) (**Fig. 5D**, yellow dots) cluster in the upper right quadrant indicating progressive accumulation in the APP-hom condition. In contrast, LRP1 and ABCG1 undergo progressive temporal suppression (**Fig. 5D**, top-left quadrant). Both proteins promote cholesterol efflux and help prevent intracellular cholesterol accumulation [48]. In the APP-het condition, cholesterol biosynthetic enzymes exhibit a significant longitudinal interaction profile; however, their absolute size effect remains tightly clustered near zero (log2 FC ∼0) (**Fig. S4A**), in contrast to their progressive accumulation in the APP-hom mutant.

To identify coordinated temporal patterns of protein expression during iNeuron maturation in the APP-London mutants, we grouped significant Interaction (Time × Genotype) proteins using K-means clustering (k=5) based on their z-score-normalized abundance profiles across weeks 2, 4, 6, and 8 (**Fig. 5E**). Of the 1,532 significant Interaction proteins, 98 targets had missing values at one or more time points in the proteomics dataset, leaving 1,439 fully quantified interaction proteins for downstream modeling. Cluster 1 (n = 365) comprised proteins that increased progressively during differentiation, whereas Cluster 2 (n = 309) comprised proteins that steadily declined over time. Trajectories in these two clusters were largely conserved across genotypes; however, APP-hom iNeurons exhibited delayed induction of Cluster 1 proteins and delayed repression of Cluster 2 proteins, evident at week 4 (**Fig. 4E**). Cluster 3 (n = 264) and 5 (n = 309) represented transient expression programs that peaked during week 4 before declining. Notably, APP-hom mutant neurons displayed delayed trajectory and higher transient expression of these protein modules, respectively. Cluster 4 (n = 192) displayed a biphasic trajectory characterized by an early decline followed by recovery during late differentiation. APP-hom mutants exhibited a more pronounced reduction at Week 4, then converged with the other genotypes at Week 8. A global heatmap tracking the 1,439 core interaction proteins into their respective K-means block complements these trajectories (**Fig. S4B**)

To determine the biological processes represented by each temporal expression program, we performed Gene Ontology, Reactome, and pathway enrichment analyses for proteins within each K-means cluster. Notably, statistical significance was dominated by cluster 2, which was strongly enriched for cholesterol and sterol biosynthesis (**Fig. 5F**). These results are consistent with previous analysis (**Fig. 5D, S3D**), indicating altered temporal accumulation of cholesterol biosynthesis in APP-hom iNeurons. Analysis also revealed additional perturbations in RNA processing, GPCR and calcium signaling, as well as synaptic maturation and neuronal connectivity (**Fig S4C**). We then aimed to dissect the temporally aberrant accumulation in cholesterol biosynthesis. We analyzed z-score trajectories for core enzymes that participate in the mevalonate-cholesterol biosynthetic pathway (**Fig. 6A**). While no significant temporal changes were evident in the cholesterol-sensing regulators SREBP2, SCAP, and INSIG2, a pronounced temporal amplification was detected for enzymes of the cholesterol biosynthetic pathway in APP-hom iNeurons. This includes enzymes in the upper mevalonate pathway (HMGCS1, HMGCR, MVK), lower mevalonate reactions (MVD, FDPS, FDFT1), and post-squalene biosynthesis (SQLE, DHR7, DHR24). The most pronounced increase occurs in week 4, particularly for the rate-limiting enzyme HMGCR (**Fig. 6A, red arrowhead**). Trajectories also indicate that expression of these enzymes peaks in early differentiation and then steadily declines, similarly to APP (**Fig. 3A**), except for HMGCR, which peaks at week 8 across genotypes. Overall, the cholesterol biosynthetic enzymes HMGCR, HMGCS1, MVK, MVD, IDI1, FDPS, FDFT1, SQLE, LSS, DHCR7, and DHCR24 exhibited a transient expression profile, peaking during week 2 of differentiation and then progressively declining (**Fig. 6B**). Similarly to the interaction results of the linear model (**Fig. 5D**), in the APP-hom condition relative and log-2 centered abundance of the cholesterol biosynthesis enzyme set showed a clear sustained upregulation during weeks 2 and 4 (**Fig. 6B-C**). However, increased expression does not necessarily indicate increased enzymatic activity.

**Figure 6.**
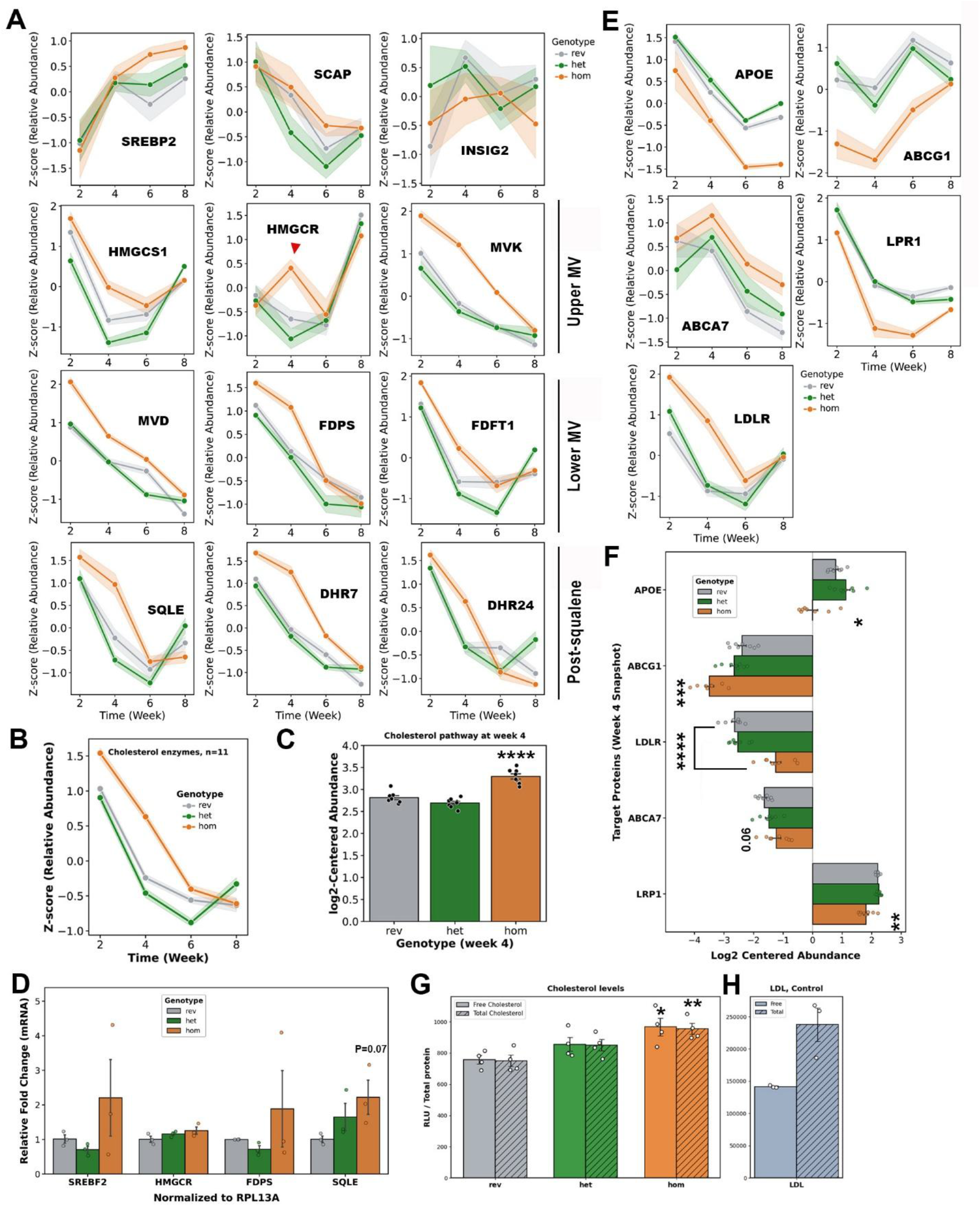
Aberrant protein upregulation in the cholesterol biosynthesis pathway and increased cholesterol accumulation in iNeurons carrying the APP-London mutation. **(A)** Longitudinal Z-score trajectories for intracellular cholesterol sensing machinery (SREBP2, SCAP, INSIG2), enzymes involved in upper mevalonate reactions (HMGCS1, HMGCR, MVK), lower mevalonate cascades (MVD, FDPS, FDFT1), and post-squalene processing (SQLE, DHCR7, DHCR24) in the APP genotypes across 8 weeks of differentiation (x-axis). **(B)** Composite line trajectories and **(C)** Log2-centered abundance bar plot at week 4; each point represents the mean log2-centered abundance of the 11 predefined cholesterol biosynthesis-associated enzymes within each sample (n=8 samples per genotype). Bars and error bars indicate the group means and SEM, respectively. Individual enzymes were not treated as independent replicates (**** P-Adj, <0.0001). **(D)** Bar plot with quantitative RT-PCR results of mRNA levels of selected genes (HMGCR, FDPS, SQLE) in APP-London iNeurons at week 4. **(E)** Protein trajectory Z-profiles of key cholesterol transporters involved in brain cholesterol homeostasis, including APOE, ABCG1, ABCA7, LRP1, and LDLR, across the three APP genotypes over time (x-axis). **(F)** Graph bars representing individual transporter abundances at week 4. Adjusted P values; *P<0.5, **P<0.01, ***P<0.001, ****P<0.0001. **(G)** Bar plot with results of free and total cholesterol levels determined by a luminescence-based assay in iNeuron lysates grown under cholesterol-free conditions for 4 weeks. Adjusted P values; *P < 0.05, **P < 0.01. **(H)** Human LDL was used as a positive control for detecting esterified cholesterol in the assay. In all trajectory plots, the shaded region and error bars in bar graphs indicate the standard error of the mean (SEM).

Transcriptional analysis of SREBP2 targets showed that mRNA fold-change values for HMGCR, FDPS, and SQLE were not significantly different (**Fig. 6D**). We also analyzed the expression of cholesterol receptors/transporters that regulate cholesterol homeostasis in the brain, including APOE, ABCG1, ABCA7, LPR1, and LDLR (**Fig. 6E**) [49]. Results showed altered expression across time, with marked dysregulation in APP-hom compared with APP-rev and -het (**Fig. 6E**). For receptors involved in the uptake of APOE-containing cholesterol, LPR1 and LDLR showed opposite responses. A significant decrease in LPR1 and an increase in LDLR were observed at week 4, with APOE abundance progressively decreasing at all time points (**Fig. 6E, F**). Interestingly, the ATP-binding cassette transporter ABCG1, which mediates cholesterol efflux, shows sustained downregulation from week 2 to 4, with striking repression at week 4 in APP-hom cultures(**Fig. 6E, F**). Consistent with a model in which positive regulation of the SREBP2 signaling pathway persists in APP-hom neurons, ABCA7 and LDLR protein abundance increase, whereas LPR1 abundance progressively decreases across time points (**Fig. 6E, F**) [50–52].

These results prompted us to examine cholesterol levels in cell lysates from APP-London iNeurons using a luminescence-based assay, in which light output is proportional to cholesterol levels in the samples. In support of a model with aberrant positive feedback on sterol regulatory element-binding protein (SREBP2)-mediated cholesterol biosynthesis concomitant with reduced cholesterol efflux, free and total cholesterol levels were significantly increased in APP-hom iNeurons (**Fig. 6G**). Of note, free and total cholesterol levels were similar in magnitude, indicating that esterified cholesterol content is negligible in iNeurons derived from the iPSC line KOLF2.1J. In parallel, low-density lipoprotein (LDL) was used as a control for esterified cholesterol levels (**Fig. 6H**), supporting low levels of esterified cholesterol in APP-London iNeurons. We cultured iNeurons in media without any source of exogenous cholesterol. Therefore, increases in cholesterol levels do not reflect alterations resulting from enhanced uptake from the medium. However, internalization of cholesterol-containing extracellular vesicles shed from iNeurons required further examination [53]. In contrast to our findings, another study using AD patient iPSC-derived neuronal cultures harboring APP duplication observed an accumulation of cholesterol esters, suggesting that gene alterations associated with familial AD can affect specific cholesterol pathways, resulting in altered cholesterol homeostasis [16].

In iNeurons matured for 4 weeks, we performed a quantitative proteomic analysis in APP-hom iNeurons following pharmacological perturbation of cholesterol production and APP amyloidogenic processing. Cholesterol-related interventions included Atorvastatin, Fatostatin, and Efavirenz [16], whereas amyloid-processing interventions included the β-secretase inhibitor IV and γ-secretase inhibitor Compound E. We applied compounds at the indicated concentrations (**Fig. 7A**) for four days, and incubated APP-hom and APP-rev cultures with vehicle DMSO (totaling 56 samples, n=8 per condition). Across the dataset, we identified approximately 8,400 proteins per sample, with a mean missing-value percentage of 16%, indicating broad and consistent proteome coverage across experimental conditions (**Supplementary Tables 6, 7, 8, and Fig. 7B-C**). We also identified a homogeneous distribution of protein intensities across the 56 samples (**Fig. 7D**). Principal component analysis (PCA) of the whole proteome did not reveal clear separation among treatments and between the two APP-London genotypes incubated with vehicle (**Fig. S5A**). As we previously observed in the longitudinal proteomic dataset, APP-hom proteomes are slightly distinct across the time points but show greater variance at week 4 (**Fig. 1H**). In addition, the lack of broad proteome-wide separation in APP-hom pharmacological-perturbation cultures suggests that a single compound dose produces discrete, specific molecular effects rather than extensive global proteome remodeling. In support, a one-way ANOVA comparing APP-hom DMSO with the tested compounds identified 161 proteins (1.82% of the quantified proteome) with a significant overall condition effect after Benjamini-Hochberg correction (FDR < 0.05; n≥ 4 observations per condition). Follow-up Welch’s t-tests comparing each compound with APP-hom DMSO identified 134 significant proteins for Atorvastatin and 4 for Compound E (**Supplementary Table 9**). Notably, no significant proteins were detected for other compounds despite their specific effects on Aβ secretion (**Fig. 7H-I**). These results indicate that Atorvastatin and Compound E show detectable compound-associated proteomic changes, with the strongest effect observed for Atorvastatin. Functional enrichment analysis showed that proteins altered by Atorvastatin were significantly enriched in cholesterol-biosynthesis pathways consistent with its established mechanism of action (**Fig. S5A**). For compound E, significantly altered proteins include upregulation of two well-established γ-secretase substrates, APLP1 and NRG1 [54,55]. APBB1, which is an interactor of the APP intracellular domain (AICD) that promotes APBB1-mediated nuclear transcription [56,57]. Unexpectedly, γ-secretase inhibition with compound E also significantly upregulated Synaptotagmin 4 (SYT4), uncovering a potential γ-secretase-associated factor that warrants further validation (**Fig. S5B**).

**Figure 7.**
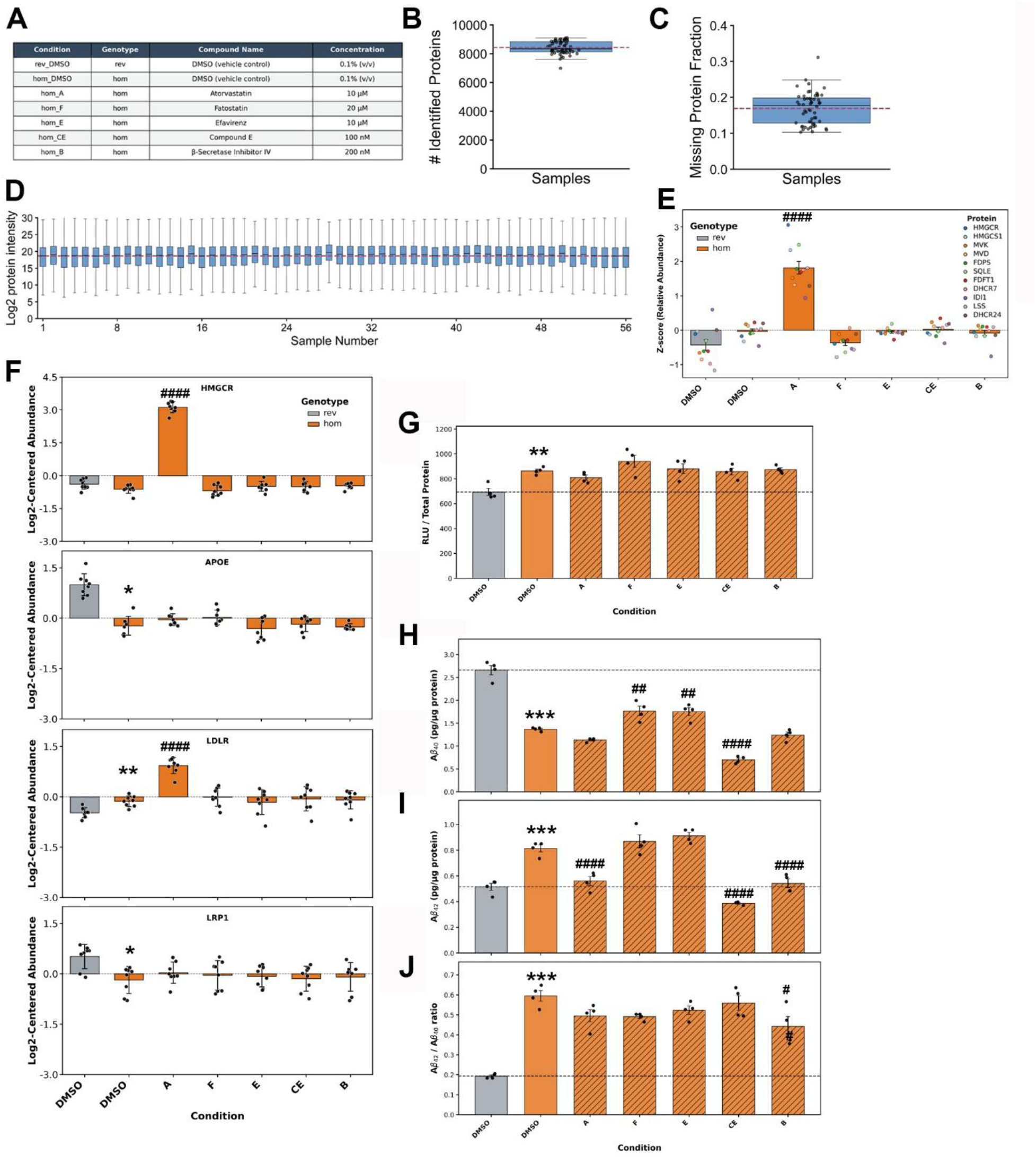
Targeted pharmacological perturbation and its effects on cholesterol metabolism and amyloid-β production. **(A)** Experimental design showing the pharmacological compounds and concentrations tested. Compounds were administered during the final four days of neuronal maturation, before completion of the week-4 endpoint. Quality-control metrics for the proteomic dataset: **(B)** number of proteins identified per sample, **(C)** fraction of missing protein-abundance measurements per sample, and **(D)** distribution of log2-transformed protein intensities. **(E)** Relative abundance Z-score for a panel of 11 enzymes associated with the sterol regulatory element-binding protein 2 (SREBP2) pathway across the indicated treatment conditions. Statistical significance was assessed using one-way ANOVA followed by Dunnett’s multiple-comparison test, comparing each compound-treated group with APP-hom DMSO. **(F)** Log2-centered abundances for specific proteins, including HMGCR, APOE, LDLR, and LRP1, shown in separate panels across treatments. Bars denote means ± SD, and overlaid individual points represent independent biological replicates. The horizontal dashed line denotes the experiment-wide centered baseline (=0.0, corresponding to the mean abundance across the complete dataset). **(G)** Total cholesterol abundance, measured after cholesterol esterase treatment to measure both free cholesterol and cholesteryl esters. **(H)** Amyloid-β (Aβ) secretion profiles across the indicated treatment conditions. **(I)** Secreted Aβ42 levels. **(J)** Aβ42/Aβ40 ratio. Asterisks indicate comparisons between APP-rev and APP-homozygous in DMSO-treated conditions. Hash symbols indicate comparisons between the APP-hom DMSO control and the indicated APP-hom pharmacological treatment. Statistical significance of adjusted P values is denoted as follows: * or # (P<0.05), ** or ## (P<0.01), *** or ### (P<0.001), and **** or #### (P<0.0001). P values were adjusted for multiple comparisons as described in the Methods section.

We next examined the protein abundance (mean z-scores) of enzymes involved in cholesterol biosynthesis (**Fig. 7E**). Under vehicle conditions, the aggregate abundance of these enzymes was slightly higher in APP-hom cultures than in APP-rev. However, this difference was not statistically significant, as we found in the longitudinal dataset (**Fig. 6 B-C**), reflecting technical variability in the proteomic datasets and a small effect size [58]. We then compared APP-hom iNeurons treated with specific compounds against APP-hom vehicle control; only Atorvastatin showed a significant increase in the relative abundance of cholesterol-biosynthesis enzymes (**Fig. 7E**, Dunnett-adjusted P< 0.0001), including its target, the rate-limiting enzyme of the pathway, HMGCR (**Fig. 7F**). Consistent with a model in which Atorvastatin increases the SREBP2 transcriptional activity, a significant increase is also detected in LDLR (**Fig. 7F**). This indicates the undesired compensatory mechanism in the SREBP2 pathway in response to statin inhibition of HMGCR [59]. In contrast, for Fatostatin, we detected no significant shifts in the protein abundance of enzymes in the mevalonate pathway, LDLR, or LRP1 relative to APP-hom DMSO (**Fig. 7E-F**), suggesting that the SREBP2 pathway is maintained at a baseline similar to the APP-rev DMSO condition [60].

Consistent with the longitudinal proteomic results, analysis of protein expression of the cholesterol transporters APOE and LDLR showed a significant decrease in APP-hom cultures compared with APP-rev cultures incubated with DMSO. This comparison also resulted in a significant decrease in the cholesterol regulator LRP1 in APP-hom iNeurons. Low protein coverage was observed for ABCG1 in this proteomic dataset, with the protein detected in only four technical replicates in the APP-hom DMSO condition (**Fig. S5B**). Although a significant decrease was detected in the longitudinal dataset for ABCG1 in APP-hom cultures, where the protein was identified in all replicates (**Fig. 6F**), we did not find evident changes in the cholesterol homeostasis machinery in cultures treated with Efavirenz, β-secretase inhibitor IV, and γ-secretase inhibitor Compound E (**Fig. 7 E-F**).

We next measured total cholesterol content in cell lysates from parallel pharmacologically perturbed cultures. Consistent with previous results (**Fig. 6G**), we observed a significant increase in APP-hom compared with APP-rev in the DMSO condition (P-Adj < 0.01) (**Fig. 7G**). Whereas none of the compounds shifted cholesterol levels in APP-hom-treated cultures, including Atorvastatin, which positively modulates mRNA and protein levels of SREBP2-target genes, or Fatostatin, which is expected to transcriptionally repress this pathway (**Fig. 7E**) [16]. Notably, LRP1 expression remained unchanged in APP-hom cultures treated with individual compounds compared to the APP-hom DMSO condition, in which expression is significantly downregulated relative to the APP-rev genotype. Overall, these suggest that alterations in LRP1 and ABCG1 cholesterol transporters are, in part, responsible for cholesterol disturbances in APP-hom cultures. However, further studies are needed to investigate the precise molecular mechanisms and the involvement of the SREBP2 axis and other pathways to explain intracellular cholesterol accumulation in APP-hom cultures [61,62].

Finally, we performed ELISA assays to assess Aβ40 and Aβ42 secretion profiles in the conditioning media. In vehicle conditions, APP-hom showed a significant increase in the Aβ42/Aβ40 ratio (**Fig. 7J**), driven by decreased Aβ40 and increased Aβ42 concentrations (**Fig. 7H and I**). In APP-hom pharmacological-perturbed cultures, atorvastatin decreased Aβ42 levels, and the Aβ42/Aβ40 ratio remained unchanged relative to APP-hom DMSO. Fatostatin and Efavirenz significantly increased Aβ40 levels while Aβ42 remained unchanged (**Fig. 7H**). As expected, γ-secretase inhibitor Compound E significantly decreased both Aβ40 and Aβ42, and the β-secretase inhibitor IV selectively decreased Aβ42 levels, resulting in a lower Aβ42/Aβ40 ratio compared to the APP-hom DMSO condition (**Fig. 7H-J**).

Results indicate that pharmacological treatments did not elicit statistically significant reductions in total cellular cholesterol compared to the vehicle control (APP-hom, DMSO). However, cholesterol was moderately and positively correlated with Aβ40 by Pearson analysis in APP-hom cultures treated with DMSO and the compounds (r=0.52, Holm-adjusted P =0.019; correction across the two prespecified Pearson tests for Aβ40 and Aβ42) (**Fig. S6C**). Cholesterol was also positively correlated with Aβ42 (r = 0.42, Holm-adjusted p = 0.042; **Fig. S5D**). Spearman correlation analysis also showed a positive association for Aβ40 (ρ=0.44, Holm-adjusted ρ=0.0644) and for Aβ42 (ρ=0.39, Holm-adjusted ρ=0.064), neither of which remained significant after correction(**Fig. S6D**). For cholesterol analysis, evaluation of individual treatment effects against the vehicle baseline (APP-hom, DMSO) revealed a narrow dynamic range of the assay readouts (Δ= −6.34 to +8.84%) (**Fig. S6C**). The concordant correlation results across treatments were largely driven by Atorvastatin (Δ cholesterol = −6.34%, Δ Aβ40 = −17.33%, Δ Aβ42 = −31.77%) and Fatostatin (Δ cholesterol = +8.84%, ΔAβ40 = +29.16%, Δ Aβ42 = +6.72%). Notably, hom-CE produced the strongest reduction in secreted Aβ40 (−48.99%) despite negligible change in intracellular cholesterol content (−0.69%). These observations suggest that in APP-hom, specific suppression of Aβ40 secretion can occur independently of modulation of bulk cholesterol content. Additional screening of cholesterol-lowering compounds or multiple-compound doses will be necessary to determine whether upstream cholesterol reduction can restore Aβ secretion profiles from APP-hom to APP-rev levels.

## DISCUSSION

We previously optimized a protocol to derive neuronal-enriched cultures from KOLF2.1J iPSCs with stable NGN2 integration and reported that monoallelic NGN2 integration favors expression of the CNS marker GPM6A over peripheral markers [20]. Here, in KOLF2.1J gene-edited derivatives with monoallelic NGN2 integration, we analyzed temporal proteomic changes in isogenic NGN2-induced cultures (iNeurons) carrying the APP-London mutation (V717I). We analyzed a differentiation time course of four time points (week 2, 4, 6, and 8) across APP^REV/WT^ and APP-specific single-nucleotide variants (APP^WT/V717I^) and (APPV^717I/V717I^) cultures referred to as APP-rev, APP-het, and APP-hom. Our analysis identified disrupted cholesterol homeostasis as a prominent phenotype associated with increased APP-London gene dosage.

Maturation time was the dominant source of proteomic variation, whereas genotype-dependent effects were most evident in APP-hom iNeurons. The results indicate that neuronal maturation in cultures is completed around week 4, and our observations are consistent with the establishment of functional neuronal glutamatergic cultures [4,19,22]. We found clear APP-genotype-associated differences in neuronal development programs, including synapse organization and axon guidance (**Fig. S3B**), that would require further exploration in APP hom mutants. Our findings indicate that molecular remodeling in iNeurons depends strongly on neuronal maturation state. The time-course proteomic analysis, pharmacological perturbation experiments, and inclusion of APP-hom iNeurons provided a more comprehensive approach to identifying molecular alterations associated with the APP-London mutation. Our approach is particularly pertinent because technical variables in differentiating iPSC neurons to model neurological disorders can introduce significant variability in proteomic datasets and lead to conclusions driven by an arbitrarily selected differentiation endpoint [58].

Analysis of APP showed that its abundance was highest during week 2 of differentiation and then declined, consistent with its roles in early neuronal development [26–28]. Analysis of ADAM family secretases and participating enzymes in the amyloidogenic pathway showed strong temporal regulation independent of genotype, emphasizing that neurodevelopmental processes can influence pathways directly relevant to AD biology [63,64]. At week 4, APP-hom neurons showed higher APP abundance than APP-rev and APP-het neurons, supported by protein-and peptide-level mass spectrometry analyses. Immunoblotting showed a higher mean FL-APP signal and nominal evidence for increased α-CTF abundance in APP-hom cultures at week 4. Neither α-CTF/FL-APP nor β-CTF/FL-APP ratios differed significantly from the revertant control (**Fig. S2C**), although the small sample size limits interpretation of these comparisons. These steady-state measurements do not establish preferential accumulation of either CTF, a selective shift in α-or β-secretase processing, or the mechanism underlying increased abundance of mutated APP. The increase in APP abundance is consistent with previous findings in patient-derived iPSC neurons carrying the APP-London mutation (V717I), in which elevated FL-APP levels were observed despite unchanged transcript levels of APP and APP secretases [31].

Proteomic alterations were strongly dependent on APP-London allele dosage. APP-het neurons showed limited proteomic alterations, whereas APP-hom neurons showed extensive changes involving proteins that participate in multiple AD-relevant pathways. Interestingly, the number of proteins changing in the same direction in both APP mutant genotypes has been implicated in Aβ and Tau pathology and glutamatergic neurotransmission, including INPPL1/SHIP2, PLXNA4, CAMK2A, and GRIA2 [42–44,65]. This directional concordance suggests that a subset of the molecular changes amplified in APP-hom cells may represent mutation-associated biology rather than nonspecific consequences of the homozygous state. Although proteomic changes were substantially greater in APP-hom than in APP-het neurons, some phenotypes in the homozygous model recapitulate features previously reported in patient-derived neurons and AD postmortem brain tissue. These include increased APP abundance, elevated Aβ42 and Aβ42/Aβ40 ratios, and upregulation of the transmembrane protein LINGO2 (**Fig. S2D**), which is involved in neurite outgrowth deficits early in development and linked to transcriptional changes involved in synaptic dysfunction and senescence [31,64]. These findings suggest that the APP-hom genotype may amplify mutation-associated phenotypes that are more subtle in the heterozygous condition and increase sensitivity for identifying AD pathways.

iNeurons represent a model that isolates early cell-autonomous responses to the pathogenic APP-London mutation, aiding understanding of complex responses in a multicellular model such as the postmortem human brain from AD patients [13,15]. Overrepresentation analysis in APP-London mutant iNeurons revealed enrichment for terms governing GPCR signaling, axon guidance, synapse assembly, calcium signaling, and early endosome, consistent with processes involved in AD pathology (**Fig. S3 and S4**) [47,64]. Among the pathways altered in APP-hom neurons, cholesterol biosynthesis and homeostasis were consistent across complementary analyses. Cholesterol-biosynthetic enzymes showed genotype-dependent temporal trajectories, with increased abundance spanning the upper and lower mevalonate pathway and post-squalene biosynthesis reactions. These changes were accompanied by altered abundance of proteins involved in cholesterol uptake and efflux, including increased LDLR and reduced LRP1, ABCG1, and APOE. Furthermore, independent experiments in APP-hom neurons matured for 4 weeks showed an increased free and total cellular cholesterol without an evident transcriptional activation of the canonical SREBP2 pathway. For instance, the SREBP2-target transcripts HMGCR, FDPS, and SQLE did not increase significantly in week-4 matured APP-hom iNeurons. This apparent uncoupling raises the possibility that the APP-London mutation affects cholesterol metabolism downstream of transcription, including changes in intracellular sterol compartmentalization or defects in endoplasmic reticulum (ER) cholesterol-sensing machinery [66,67].

In APP-hom neurons, increased total cholesterol accompanied increased APP and HMGCR abundance. Previous studies suggest that APP-CTFs can disrupt cholesterol homeostasis, providing a potential mechanism to investigate in this model. This convergence is consistent with CTFs perturbing intracellular cholesterol homeostasis in an AD model. In a non-neuronal model, isogenic homozygous APP-V717I iPSCs, C99 (β-CTF) accumulation was accompanied by enhanced cholesterol uptake and esterification, and C99 was proposed to act as a cholesterol -sensing peptide that promotes cholesterol trafficking to mitochondria-associated ER membranes (MAMs) [68]. More recently, excess in APP-CTFs were shown to accumulate at late-endosome/lysosome-ER contact sites, impair lysosomal Ca²⁺ refilling, promote cholesterol accretion and subsequently disrupt endosomal recycling and maturation [69]. Consistent with a broader connection between APP-CTFs and endosomal homeostasis, β-CTF accumulation in isogenic APP-and PSEN1-mutant human neurons has been associated with Rab5-positive early-endosome enlargement independently of Aβ [47]. These observations suggest plausible pathways through which increased APP catabolic products could influence both cholesterol distribution and endolysosomal function.

In our model, however, although the absolute abundance of APP CTFs increased, neither α-CTF/FL-APP nor β-CTF/FL-APP ratios differed significantly from the revertant control. Thus, our results indicate an increased steady-state burden of APP-derived species without evidence of preferential enhancement of α-or β-secretase processing. Our findings therefore establish concurrent changes in APP abundance and cholesterol homeostasis but do not determine whether CTF abundance contributes directly to the cholesterol phenotype and whether these fragments accumulate within specific endosomal or ER membrane compartments.

Changes in cholesterol transport proteins further suggest that impaired cholesterol redistribution and/or flux may contribute to the phenotype in APP-hom cultures. LRP1, ABCG1, and APOE were reduced, whereas LDLR increased. Although protein abundance alone cannot establish flux, the combination of increased biosynthetic enzymes, altered transporter expression, and elevated intracellular cholesterol is consistent with disruption at multiple levels of cholesterol homeostasis [68,70]. Reduced LRP1 is particularly notable because APP processing may provide a transcriptional link between APP and lipoprotein metabolism. APP intracellular domain (AICD), along with Fe65 and Tip60, has been reported to associate with the LRP1 promoter and suppress LRP1 transcription, thereby influencing APOE expression and cholesterol metabolism [71]. Thus, it is possible this pathway may be contributing to altered LRP1 abundance in APP-hom iNeurons.

The predominance of free rather than esterified cholesterol in this model is also notable. Previous studies in other familial AD neuronal models have described cholesteryl ester accumulation [16,68], suggesting that distinct APP alterations may perturb distinct cholesterol metabolism pathways. Moreover, pathogenic alterations in γ-secretase activity have previously been linked to reciprocal changes in cholesterol and sphingomyelin metabolism, further emphasizing that changes in lipid composition may extend beyond total cholesterol abundance [72]. Further studies defining the subcellular localization and metabolic fate of accumulated cholesterol would clarify the physiological implications of the reported disturbances observed in APP-V717I iNeurons.

Our pharmacological perturbation experiments further show that the relationship between APP-London and neuronal cholesterol homeostasis is not explained solely by biosynthetic enzyme abundance or SREBP2 transcriptional activity. Atorvastatin induced a significant compensatory increase in the rate-limiting enzyme HMGCR and cholesterol-biosynthetic enzymes associated with sterol feedback regulation. In contrast, Fatostatin did not substantially alter the cholesterol-biosynthetic protein profile [59]. Despite these molecular responses, these statins did not reduce the elevated cholesterol levels in APP-hom cultures. It is possible that drug treatments required further dose optimization to produced a detectable difference in cholesterol phenotypes. None of the tested compounds decreased cholesterol to the levels detected in APP-rev cultures. Interestingly, proteomics analysis consistently identified reductions in APOE and LRP1 alongside an increase in LDLR in APP-hom cultures. These observations imply that the altered cholesterol phenotype involves multiple pathways that may require a combination of drugs to reestablish cholesterol disturbances.

Our results also argue against a simple linear relationship between total intracellular cholesterol and Aβ production. APP-hom neurons showed the expected shift toward an increased Aβ42/Aβ40 ratio under vehicle conditions, whereas pharmacological interventions produced distinct effects on Aβ40 and Aβ42 without changing total cellular cholesterol. Although cholesterol showed a moderate positive association with both Aβ40 Aβ42across treatment conditions, this relationship varied by compound and did not remain significant for either Aβ40 or Aβ42 after multiple-testing correction (**Fig. S6C-D**). Most notably, γ-secretase inhibition markedly reduced secreted Aβ40 and Aβ42 as expected despite unchanged intracellular cholesterol. Thus, APP amyloidogenic processing can change independently of detectable changes in bulk cellular cholesterol. Cholesterol and APP processing are therefore likely to intersect at specific subcellular compartments rather than through a simple relationship between total cellular cholesterol concentration and Aβ secretion [73]. Unexpectedly, pharmacological inhibition of γ-secretase with compound E significantly upregulated Synaptotagmin 4 (SYT4). Synaptotagmin isoform 7 (SYT7) expression and localization at the plasma membrane are regulated by γ-secretase activity [74,75]. Moreover, both SYT4 and SYT7 are required for neurotransmitter release at the somatodendritic compartment [76]. Taken together, this suggests SYT4 as a potential γ-secretase-associated factor and/or substrate that warrants further validation (**Fig. S5B**).

Overall, our findings identify a maturation-dependent molecular phenotype of APP-London neurons, characterized by increased APP abundance at week 4 and disrupted cholesterol homeostasis in the homozygous genotype. The increase in abundance of cholesterol biosynthetic enzymes, altered expression of cholesterol transport machinery, and elevated intracellular cholesterol suggest that lipid dysregulation is an early molecular event in response to the APP-London mutation. In addition, the absence of concordant SREBP2-driven transcriptional activation and the limited ability of cholesterol-directed pharmacological interventions to reduce intracellular cholesterol argue for a more complex regulatory mechanism than simple activation of the SREBP2 pathway [15]. These results will therefore motivate investigation of post-transcriptional regulation, cholesterol trafficking and efflux, and compartment-specific sterol sensing as potential mechanisms linking the pathogenic APP-London mutation to neuronal lipid dyshomeostasis and amyloid biology.

Limitations of the study: The proteomic experiments were performed in a well-characterized iPSC line [18]; however, to establish generalizability of findings, independent genetic backgrounds may be required. The reported shifts in protein abundance do not necessarily reflect enzymatic activity or function, and the present experiments do not directly measure cholesterol synthesis or flux. Biochemical measurements of total and free cholesterol cannot resolve potentially disease-relevant differences among cholesterol pools in different cellular compartments. Finally, the pharmacological experiment evaluated a limited set of compounds and conditions, and compensatory feedback may obscure the effects of short-term pathway inhibition.

To support broader use of this longitudinal dataset, we developed an interactive web resource that lets users explore the temporal abundance of profiles of individual proteins across APP-London genotypes and stages of neuronal maturation. The streamlit application is available free here https://neurons-timecourse-app.streamlit.app/. This resource is intended to provide a complementary, disease-focused resource for investigators using KOLF2.1J-based neurodegeneration models.

## METHODS

### Development and culturing of iNeurons for proteomic analysis

Human-induced pluripotent stem cells (iPSCs) harboring the London APP mutation (V717I) were obtained from The Jackson Laboratories. iPSC cultures were maintained and propagated according to the supplier’s instructions. Generation KOLF2.1J-derived genome-edited iPSC lines carrying a doxycycline-inducible NGN2 cassette, as well as the neuronal induction strategy, were previously described [19]. NGN2 integration at the AAVS1 site was confirmed by PCR-based genotyping using validated primers [4], and in NGN2 stable iPSC lines, G-banding karyotyping at a resolution of 350-375 was performed by WiCell Research Institute (Fig. S1A).

Briefly, iPSCs were dissociated to single cells using Accutase, and approximately 1.8 x 10^6 cells were plated onto Matrigel-coated 10-cm dishes in DMEM/F12 supplemented with N2, GlutaMAX (1X), NEAA (1X), the ROCK inhibitor Y-27632, and doxycycline (2 µg/mL). After 24 h, the medium was replaced with fresh induction medium containing doxycycline but without the ROCK inhibitor. Then, cells were maintained for 48 h with daily media changes. The resulting enriched-neuronal progenitor populations were subsequently dissociated with Accutase and cryopreserved in liquid nitrogen at 1.2 x 10^6 cells per vial until use. Prior to plating at each time point, cells from vials stored in liquid nitrogen were recovered, and cell viability was assessed by trypan blue staining, with consistently >90% viability. 1 x 10^5 cells were plated into individual wells of poly-L-ornithine (PLO)-coated 96-well plates in BrainPhys Neuronal Medium supplemented with BDNF (10 ng/mL), NT-3 (10 ng/mL), B27 (1X), mouse laminin (1 µg/mL), and doxycycline (3 µg/mL) and DAPT (5 µM) for 24 h. Cultures were rinsed with fresh medium to remove doxycycline and DAPT, and cells were maintained with half-media changes twice weekly during the neuronal maturation period.

### Proteomic sample preparation and analysis

For the three APP-London genotypes, iNeuron cell lysates were collected for each time point (2, 4, 6, or 8 weeks) (n=8 technical replicates per condition, totaling 96 samples). Cells were rinsed with PBS and lysed in RIPA buffer (ThermoScientific, #J62524.AE) supplemented with protease inhibitors. Proteomic sample preparation for bulk proteomic analysis was performed as described (PMID: 42309337). Briefly, protein concentration of iNeuron lysates was determined using the BCA (Pierce) assay. Approximately 20 µg of protein per sample was processed using a modified SP3 workflow (PMID: 30464214). Protein lysates were subjected to reduction and alkylation (DTT/IAA), captured on silica magnetic beads to remove LC-MS-incompatible chemicals, and digested overnight at 37 °C with trypsin (Promega, #V511C). Trypsin was used at a 1:30 (v/v) ratio in iNeuron protein lysates. Peptides were acidified to quench digestion and diluted prior to LC-MS/MS analysis on an Orbitrap Astral mass spectrometer (Thermo Fisher Scientific) in data-independent acquisition (DIA) mode. Peptides were separated on a PepSep C18 analytical column using a 30-min gradient, with randomized injection order and periodic injections of HeLa digest to assess optimal instrument performance.

Data were processed using DIA-NN (v2.2.0) software against the UniProt database (UP000005640, downloaded 12/01/2025). The main search was against the human Uniprot reference proteome (Uniprot accession UP000005640, downloaded on 12/01/25), which contains 20,632 proteins. Protein identification was performed with Trypsin/P specificity, with up to one missed cleavage, and N-terminal methionine excision was enabled. Peptide length was restricted to 7-30 amino acids, and precursor charge states to 1-4, precursor m/z values of 300-1800, and fragment ion m/z values of 200-1800 were considered. Deep learning-based prediction of fragment ion spectra, retention time, and ion mobility was enabled for library generation. Match-between-runs (MBR), RT-dependent cross-run normalization, and RT/ion mobility profiling were enabled. Heuristic protein inference was applied, and AuantUMS (high precision) was applied as a quantification strategy. RT-dependent cross-run normalization; library generation depends on IDs, RT, and IM profiling. Precursor-and protein-level identifications were controlled at a 1% false discovery rate (FDR) using the built-in DIA-NN q-value–based filtering algorithm.

### Proteomic data preprocessing and computational analysis

Protein abundance values in the longitudinal dataset were log2-transformed, and protein counts that deviated by more than ±15% from the dataset mean were excluded prior to downstream analysis. Of 96 samples, 3 were excluded. Sample-to-sample correlation analysis was performed on all retained samples using their log2-transformed and median-centered protein abundance profiles, which were subsequently standardized per protein as row-wise z-scores. This transformation standardized the scores to a mean of 0 and a unit variance of 1. Pearson correlation coefficients were calculated using the z-score-transformed protein abundance matrix. Technical missingness in the matrix was resolved with the K-Nearest Neighbors (KNN) imputation algorithm (k=5 neighbors) via the scikit-learn library package. To map global protein variation, Principal Component Analysis (PCA) was performed on the transposed standardized protein abundance matrix across the iNeuron maturation times (weeks) and London-APP genotype status (rev, het, and hom). To map maturation trajectories across the 8-week window, pathway-level scores were calculated for the Gene Ontology (GO) terms: Neuron Maturation (GO:0042551), Axon Development (GO:0061564), and Dendrite Extension (GO:0097484). The analysis used standardized z-scores for individual proteins, and computations were performed only for proteins with at least 75% presence in each experimental condition (n≥ 6 replicates). Heatmaps for molecular characterization of cellular lineages and selected GO pathways across the longitudinal proteomic dataset were generated using seaborn and matplotlib libraries in Python. To track the longitudinal expression profiles of selected proteins, z-scores were evaluated at each time point (2, 4, 6, and 8 weeks). For individual z-score trajectory plots across time points, protein presence was detected in 7-8 technical replicates for each APP-London genotype, except for INSIG2 and ADAM17, for which a lower identification was obtained from 3-7 technical replicates across the three genotypes.

### Longitudinal differential protein abundance analysis

To assess proteomic expression profiles across weeks 2, 4, 6, and 8 of the three distinct APP-London genotypes (rev, het, and hom), a linear model was implemented using statsmodels (Python). Prior to statistical analysis, protein abundances were log2-transformed and median-centered. To ensure robust statistical interference, only proteins quantified in a minimum of 4 technical replicates in every experimental condition (genotype and time point). After filtering, 7,319 of 9,315 quantified proteins were retained for statistical analysis. For each retained protein, an ordinary least squares (OLS) linear model was performed using the formula: protein intensity ∼ genotype x timepoint, where genotype (APP-rev, APP-het, and APP-hom) and developmental timepoints (Weeks 2, 4, 6, and 8) were treated as categorical variables. APP-rev served as the reference genotype. Overall effects of genotype, time point, and the genotype x time point interaction were evaluated using a Type-II Analysis of Variance (ANOVA). Baseline genotype contrasts (APP-het vs APP-rev and APP-hom vs APP-rev) were estimated from the linear model using Wald tests. To obtain genotype effects representative of the entire experimental time course, interaction coefficients were averaged across all developmental time points before testing. Estimated coefficients were reported as log2 fold changes (log2FC), along with their associated p-values. To account for multiple parallel hypotheses, raw p-values were corrected column-by-column across all processed proteins using the Benjamini-Hochberg (BH) False Discovery Rate (FDR) procedure. Proteins with an FDR-adjusted *P* value < 0.05 were considered statistically significant.

For targeted abundance analysis at week 4, we log2-transformed each protein’s intensity by subtracting its mean log2 abundance across all retained samples (Proteins must be present in the 8 technical replicates). We calculated a sample-level pathway score as the arithmetic mean of the log2-centered abundances, yielding one pathway score per sample. Pathways related to APP processing and cholesterol biosynthesis were included to complement the linear modeling results. We evaluated overall differences among the three APP-London genotypes using Welch’s one-way analysis of variance, which does not assume equal variances among groups. Two prespecified pairwise comparisons, APP-het versus APP-rev and APP-hom versus APP-rev, were subsequently evaluated using two-sided Welch’s independent-samples t-tests. Pairwise P values were adjusted across the two planned contrasts using the Holm procedure to control the family-wise error rate. Holm-adjusted (P<0.05) was considered statistically significant.

To evaluate APP abundance at the peptide level, duplicated peptide sequences were averaged within each sample (n=10 duplicated, n=25 unique sequences), and low-coverage samples and peptides with greater than 50% missingness were excluded (n=23 sequences passed QC). Genotype-specific peptide estimates were required to contain at least four out of the eight detected technical replicates (n=94 out of the 96 samples). Intensities from the retained peptides were log2-transformed and standardized independently across all retained samples to account for peptide-specific differences in baseline intensity.

### Western blot analysis

After 28 days of maturation, iNeurons were washed with PBS and lysed on ice with RIPA buffer containing protease inhibitors (Roche). Micro BCA protein kit (Thermo Scientific) was used on iNeuron lysates to determine protein concentration (Pierce). Samples were prepared in Laemmli SDS sample buffer containing β-mercaptoethanol reducing agent and heated at 95°C for 5 min prior to electrophoresis. Approximately 20 µg of protein per lane was resolved on 4-15% TGX stain-free gels (Bio-Rad) and transferred to PVDF membranes (0.22 μm) using Trans-Blot Turbo transfer system (25V/3min) (Bio-Rad). Total protein per lane was visualized using a stain-free imaging system to confirm uniform loading and enable normalization of total protein (ChemiDoc MP, Bio-Rad). Membranes were blocked in 5% nonfat milk in TBS-T (Tris-buffered saline, 0.1% Tween-20) for 1 h at room temperature and incubated with primary antibodies against Amyloid Precursor Protein (APP Y188; Abcam, #ab32136, 1:10,000 dilution) and HRP anti-GAPDH (BioLegend, #FF26A, 1:5,000 dilution) overnight at 4°C. To detect APP, after washing with TBS-T, membranes were incubated with HRP-conjugated secondary antibody for 1 h at room temperature (#5220-0335, SeraCare). Bands were detected using Amersham ECL Select™ Western Blotting detection reagent (Cytiva, RPN2235) and imaged with a ChemiDoc MP imaging system (Bio-Rad). Band intensities were quantified using Image Lab software (v 6.1). APP signals were normalized to total protein per lane (stain-free). Normalized values are reported in arbitrary units (a.u.). For each band (full-length APP (FL), α-CTF, and β-CTF), we analyzed genotype differences independently using one-way analysis of variance (ANOVA), with APP-rev as the respecified control. We performed post-hoc comparisons using Dunnett’s multiple-comparison test, with n = 3 independent biological samples per genotype. P-values from analyses of the three protein species were adjusted using the Holm method. Statistical tests were two-sided, and P < 0.05 was considered statistically significant.

### Cholesterol measurement by luminescence assay

Approximately 1 × 10^5 NGN2-induced iPSCs were seeded into individual PLO-coated wells of a 96-well plate. After 28 days (28 DIV) of maturation, cellular cholesterol was quantified using the Cholesterol/Cholesterol Ester-Glo™ luminescent assay (Promega) according to the manufacturer’s instructions. Briefly, after treatment, the culture medium was removed, and the cells were washed twice with PBS. 50 μL of the kit’s lysis buffer was added to the wells containing cells, and the plate was incubated at 37 °C for 30 minutes. Cell lysate was diluted with the kit’s cholesterol lysis buffer (1:5 dilution), and 50 μL of the diluted cell lysate was transferred into individual wells of a white-walled 96-well plate (assay plate). The assay plate also contained 50 μL of cholesterol standard dilutions prepared in lysis solution (ranging from 0 to 80 µM). Then, 50 μL of the cholesterol detection reagent was added to the wells tested in the assay plate, and the samples were incubated for 1 hour at room temperature on a plate shaker. The kit’s detection reagent is enzymatically coupled with cholesterol to generate NADH, which helps reduce a pro-luciferin reductase substrate to produce luciferin. Luciferin is detected in a luciferase reaction using UltraGlo Luciferase, and the amount of light produced is proportional to the amount of cholesterol in the sample. Luminescence was recorded using a BioTek Cytation 1 cell imaging multi-mode reader (Agilent) with an integration time of 1.0 s. Data were recorded as raw relative luminescence units (RLUs). As a positive control for cholesterol content, purified human low-density lipoprotein (LDL, 6.53mg/mL, Merck Millipore) was used and diluted in 50 µL of the kit’s lysis buffer (1:500 dilution).

### Quantitative Real-Time PCR (qRT-PCR) Analysis of SREBP2

After 72 hours of DOX treatment (NGN2 initial induction) of single iPSC clones, 1 × 10^5 cells were replated into 96-well plates and treated with DOX, DOX + DAPT (5 μM), or vehicle (control) for an additional 24 hours. We employed two cell lines with homozygous and heterozygous NGN2 integration at the AAVS1 locus (Clone 1 and Clone 5). After treatments, total RNA was isolated using RNeasy spin columns (Quiagen) according to the manufacturer’s instructions. Isolated RNA was then converted into cDNA using a cDNA synthesis kit (Thermo Fisher Scientic, Verso). Expression levels from both cell lines (n=4 per treatment condition) were analyzed using SYBR Green master mix (Bio-Rad) and gene-specific qPCR primers on a CFX96 Real-Time PCR machine (Bio-Rad). Primers for HES5, NOTCH1, NGN2, DLL1, and RPL13A primers were purchased from Qiagen (catalogue numbers: QT00202699, QT00231056, QT00020447, QT00057631, QT00089915).

Analysis of melting curves indicated amplification of a single product ensuring primer specificity. The 2^(−ΔΔCt) formula was used to calculate fold changes and RPL13A served as the reference gene. For each sample, ΔCt was first obtained by subtracting the Ct of RPL13A from the Ct of the target gene. ΔΔCt values were then determined by comparing each condition to the control group. Fold-change values represent 2^(−ΔΔCt) and reflect relative gene expression normalized to RPL13A and expression relative to cells cultured without doxycycline (or vehicle control condition).

### Amyloid-β (Aβ40 and Aβ42) ELISAs

iNeurons were cultured and maintained for 4 weeks post-NGN2 induction in 96-well plates, and conditioned media was collected from the different APP-London cultures. Collected conditioned media was supplemented with the serine protease inhibitor 4-(2-aminoethyl)benzenesulfonyl fluoride (AEBSF, 1 mM final concentration) to prevent peptide degradation, and kept on ice or stored at −80°C until analysis. Secreted human Aβ40 and Aβ42 levels were quantified using solid-phase sandwich ELISA kits (Invitrogen/Thermo Fisher Scientific: Human Aβ40, Catalog No. KHB3481; Human Aβ42, Cat. No. KHB3441) according to the manufacturer’s protocols. Optical density was measured at 450 nm on a microplate absorbance reader. Concentrations were calculated using a four-parameter logistic (4PL) standard curve fit and adjusted for dilution factors. Final Aβ peptide concentrations were normalized to total intracellular protein content determined with the BCA assay, as indicated. In Fig. 4, undiluted media was used for detecting Aβ40 levels, and in Figure 7, conditioned media was diluted (1:3).

## Data and code availability

Mass spectrometry raw data for both the Time Course Analysis and the Perturbation Drug Screen are available on massive.ucsd.edu as MSV103499. For the Time Course Analysis, Supplementary Tables 4, 5, and 6 include the DIA-NN output file, metadata, and processed data (log2, log2-centered, and z-scores for all proteins identified), respectively. For the Perturbation Drug Screen, Tables 7, 8, and 9 include the DIA-NN output file, metadata, and processed data (log2-centered, and z-scores), respectively.

All data processing code is available as Jupyter notebooks on Github https://github.com/xomicsdatascience/temporal-proteomics-APP-london-mutation

The interactive data explorer is available from https://neurons-timecourse-app.streamlit.app/ with the underlying streamlit application code available from https://github.com/xomicsdatascience/neurons-timecourse-streamlit.

## Supplemental tables referenced in the manuscript

Supplementary Table 1. Time course proteomics metadata

Supplementary Table 2. Time course proteomics raw quantities from DIA-NN

Supplementary Table 3. Time course proteomics processed protein quantities

Supplementary Table 4. Statistical Results from the Linear Model

Supplementary Table 5. Genotype Statistical Concordance Results comparing APP-hom and APP-het London mutants

Supplementary Table 6. Drug perturbation experiment metadata

Supplementary Table 7. Drug perturbation proteomics raw quantities from DIA-NN

Supplementary Table 8. Drug perturbation proteomics processed quantities

Supplementary Table 9. Drug Perturbation Experiment Statistical Results

## Acknowledgements

This work was supported by the NIH (R21AG074234). We thank Lahiruni Halwatera for collecting the temporal proteomics data.

## Author Contributions

J. M.-E. and J. G. M. conceptualization; J. M.-E., Y.J., and J. G. M. methodology; J. M.-E., J. G. M. and Y.J. software; J. M.-E. and J. G. M. formal analysis; J. M.-E. and J. G. M. investigation; J. M.-E. validation; J. M.-E. and J. G. M. resources; J. M.-E. and J. G. M. data curation; J. M.-E. writing–original draft; J. M.-E. and J. G. M. writing–reviewing and editing; J. M.-E. and J. G. M. visualization; J. G. M. supervision; J. M.-E. and J. G. M. project administration; J. G. M. funding acquisition.

