## Supplementary Figures for "Temporal Proteomics Links the APP London Mutation to Early Cholesterol Remodeling Cell Programs in Human Neurons"

**Commented [JM1]:** Trying to think of a title that covers the main discovery more clearly. Alternative:

Longitudinal Proteomics of Isogenic *APP* (V717I) Human Neurons Uncovers Post-Transcriptional Upregulation of Cholesterol Biosynthesis and Impaired Sterol Efflux

Longitudinal Proteomics of *APP* (V717I) Human Neurons Reveals Coordinated Accumulation of Mevalonate Enzymes and Transporter Remodeling Independent of SREBP2 Transcription

**Commented [JM2R1]:** I prefer not to include SREBP2 transcription in the title, we measured only a few targets of the pathway and analyzed one time point (week4).

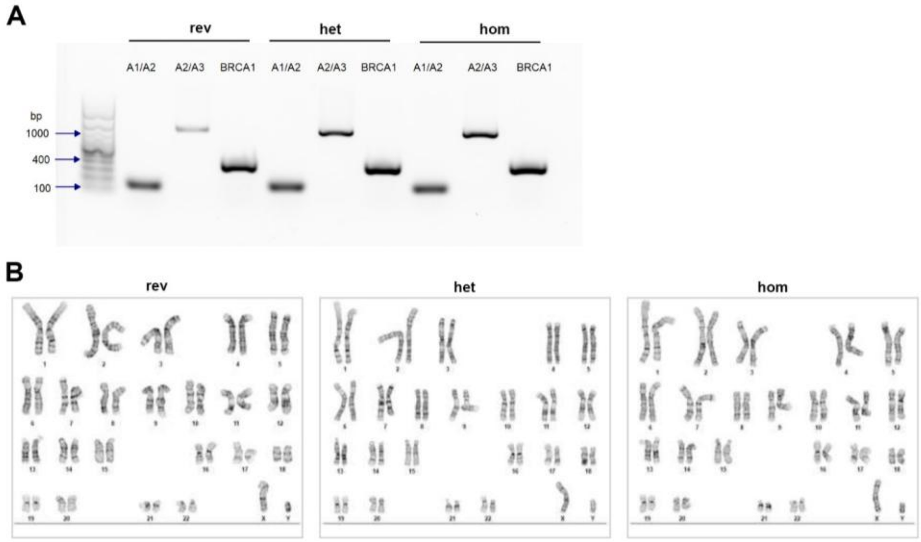


**Figure S1. Characterization of NGN2 transgene integration at the AAVS1 locus in KOLF2.1J-derived lines harboring the APP-London mutation. (A)** PCR-based genotyping in APP-rev, APP-het, and APP-hom using A1/A2 and A2/A3 primer sets [4]. BRCA1 was used as a reference gene with the following primers: F: 5'-GGTGTTTGCTACATAATGCCATATT-3', R: 5'-TTTGTTGACCCTTTCTGTTGAAG-3'. The genotype indicates monoallelic NGN2 integration in the three isogenic lines**. (B)** Representative G-banded karyotype analysis of APP-rev, APP-het, and APP-hom at a resolution of 350-375. Analysis was performed in 20 cells per genotype by WiCell Research Institute.

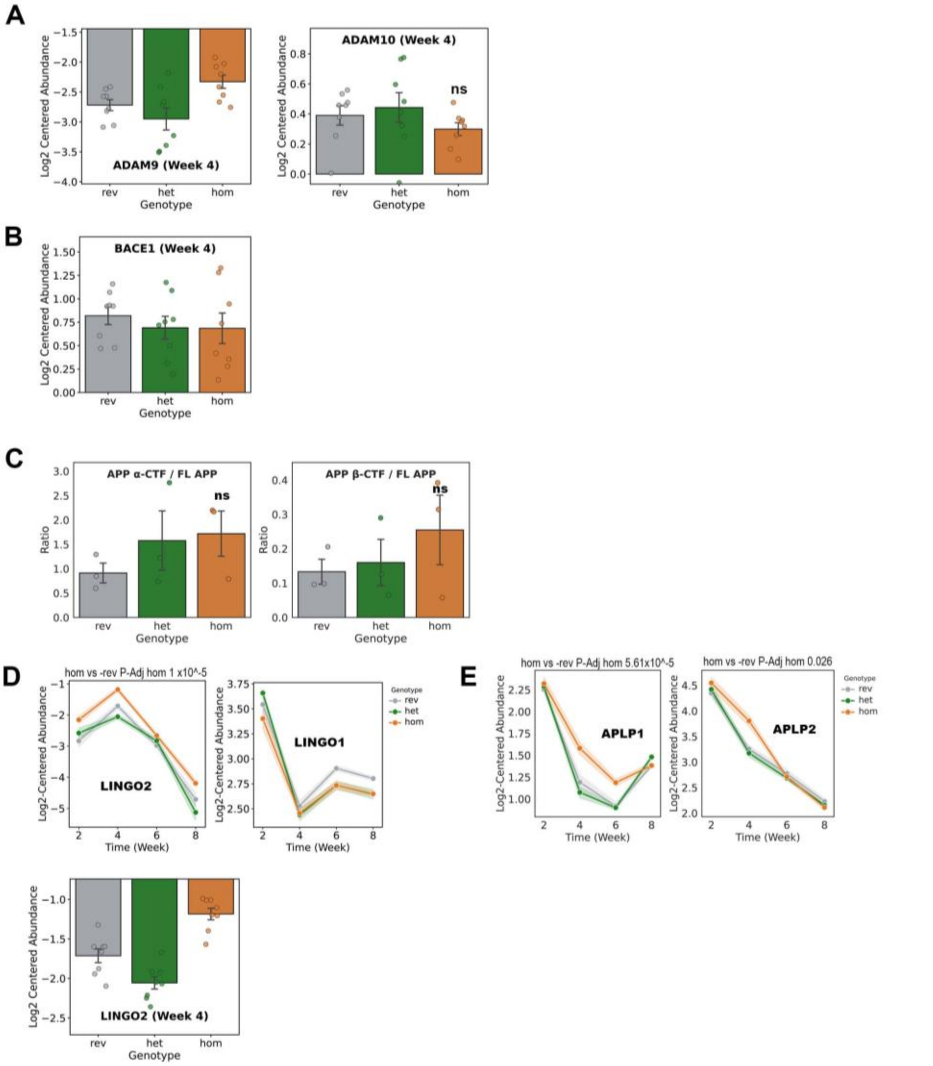


**Figure S2. Analysis of secretase abundances and APP metabolic products in iNeurons harboring the APP-London mutation. (A)** Bar plots showing log2-centered abundance (y-axis) for ADAM9 and ADAM10 across the three APP-London genotypes (x-axis) at week 4. Individual points represent biological replicates (n≥7). **(B)** Bar plot with Log2-centered abundance for the major β-secretase enzyme BACE1 at week 4 in the three APP genotypes.

**(C)** Quantification of α-CTF/FL APP and β-CTF/FL APP ratios in the three APP genotypes. Analysis associated with Western blot results in Fig. 3C. Bars represent mean ± SEM, and individual points represent independent biological samples (n=3 per genotype). We analyzed genotype effects using one-way ANOVA followed by Dunnett's multiple-comparison test, with Holm correction across comparisons. APP-rev was used as the control genotype; ns, not significant. **(D)** Longitudinal z-score trajectories for the membrane proteins LINGO1 and LINGO2 across 8 weeks of differentiation (x-axis) and a bar plot showing log2-centered abundance for LINGO1 at week 4. **(E)** Trajectories are also shown for the two APP gene family members APLP1 and APLP2 proteins across the time course; P-adj values are displayed for each one, derived from APP-rev vs -het and -hom contrasts of the linear model.

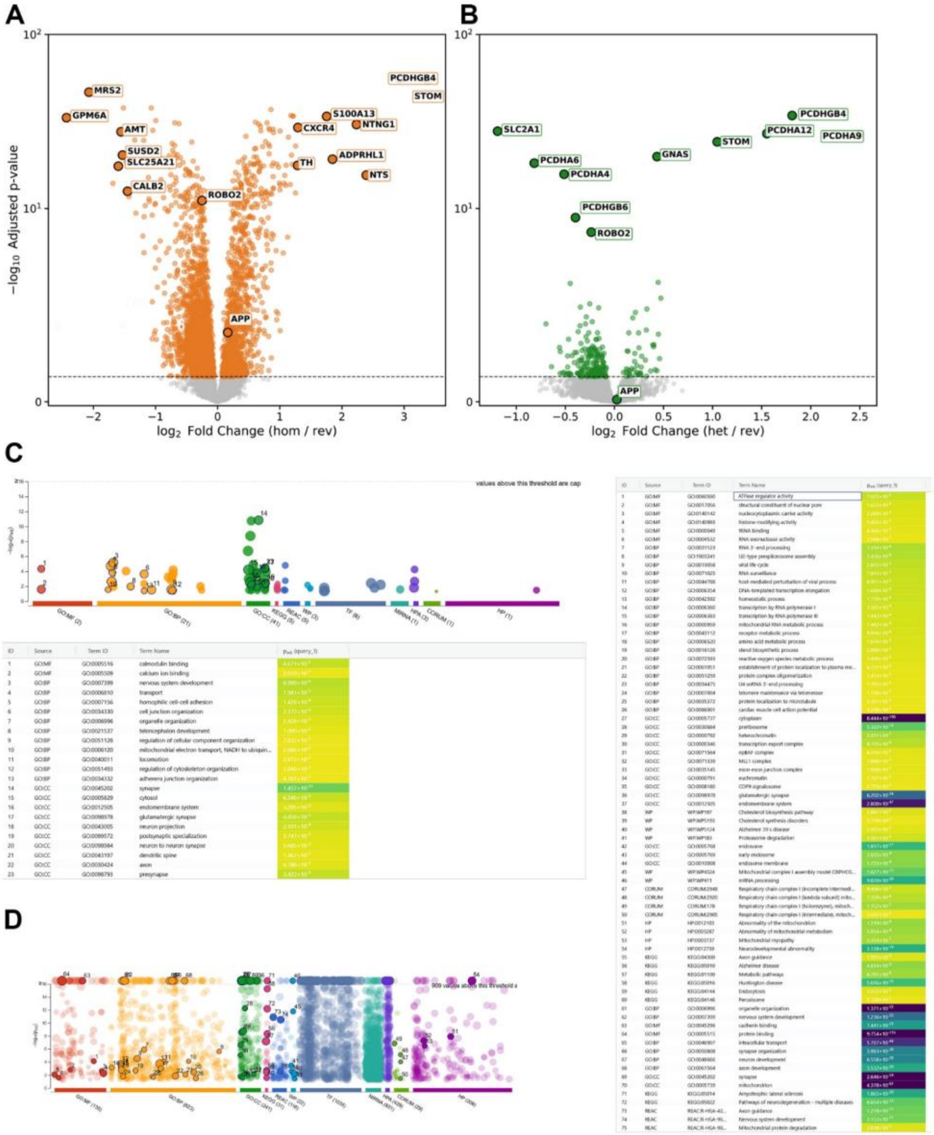

**Figure S3. Differential protein expression analysis of iNeurons carrying the APP-London mutation.** Volcano plots show differential protein abundance identified by the employed linear model for contrasts (APP-hom vs APP-rev) **(A)** and (APP-het vs APP-rev) **(B).** The x-axis represents the log2 fold change (log2 FC) relative to the APP-rev condition, and the y-axis represents −log10 of the P-Adj value. The horizontal dashed line indicates the significance level of P-Adj = 0.05. Each point represents a single protein, and annotated proteins emphasize proteins with large absolute log2 fold changes. Functional enrichment analysis of proteins exhibiting differential abundance in both APP-het **(C)** , and APP-hom **(D)** relative to APP-rev cultures. Bubble plots show g: Profiler functional enrichment of significant proteins. The y-axis represents enrichment significance (-log10 P-Adj value), and the x-axis lists the significant terms grouped by data sources. In (C) and (D), detailed tables list significant term identifications and P-Adj values.

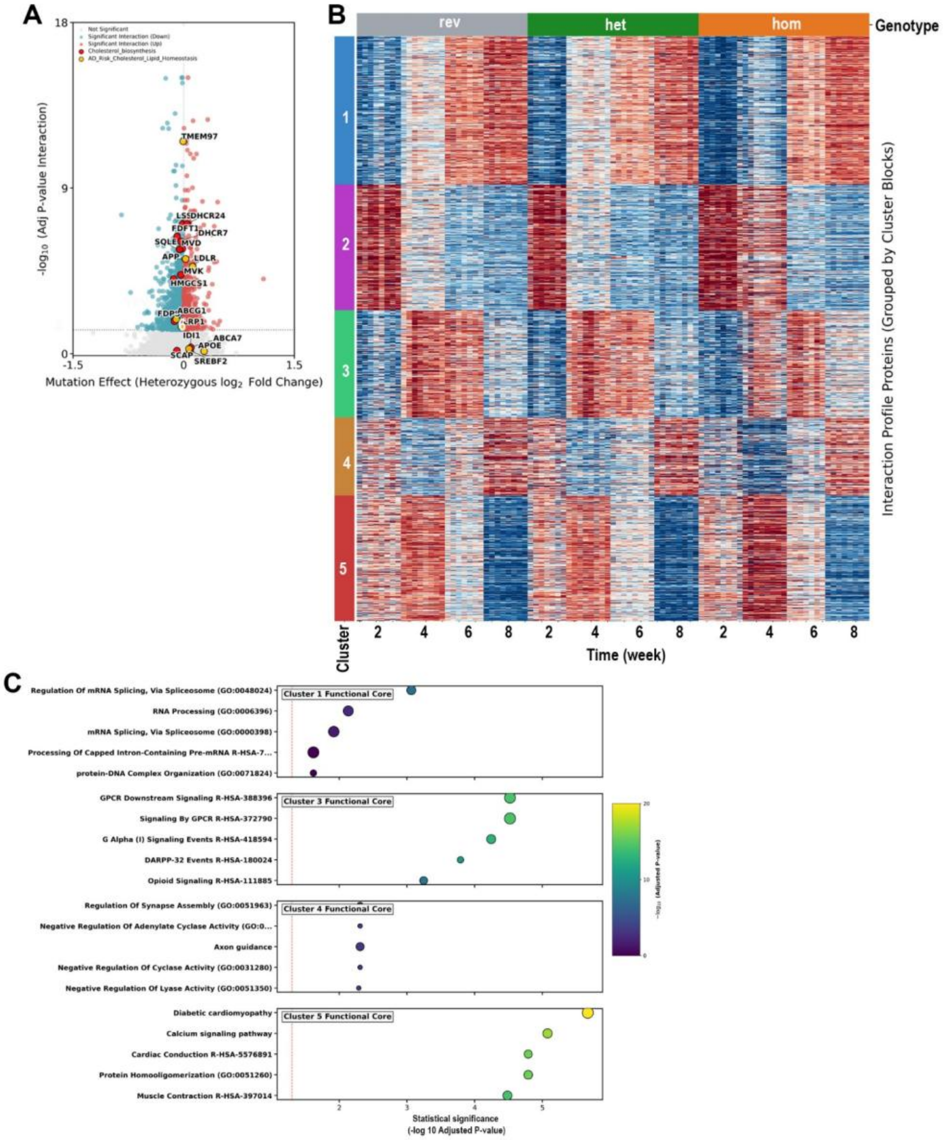


**Figure S4. Linear model results of the longitudinal proteomic dataset from iNeurons harboring the APP-London mutation. (A)** Volcano plot showing the APP-het Genotype effect (APP-het vs. APP-rev log2 fold change; x-axis) and the significance of the interaction

Time x Genotype (−log10 P-Adj value; y-axis). The vertical dotted line indicates no change relative to the APP-rev group (log2FC = 0), and the horizontal dotted line denotes the significance threshold (P-Adj = 0.05). Significant proteins are shown in red (positive log2FC) or blue (negative log2FC), whereas nonsignificant proteins are shown in gray. Proteins involved in cholesterol biosynthesis are colored red, and proteins involved in cholesterol transport are highlighted in yellow. **(B)** Heatmap of proteins identified by the linear modeling that exhibited significant interactions, grouped by K-means clustering. Protein abundances were standardized as row-wise Z-scores prior to K-means clustering (K = 5). Columns represent individual samples in APP-rev, APP-het, and APP-hom iNeurons across maturation time points (2, 4, 6, and 8 weeks), and the colored bar on the left denotes cluster assignment. Heatmap colors indicate relative protein abundance (red, higher abundance; blue, lower abundance) after row-wise z-score normalization. **(C)** Bubble plot displaying significantly enriched biological terms with proteins assigned to Clusters 1, 3, 4, and 5. Top 5 enriched terms (y-axis) with significance (x-axis). Bubble color reflects P-Adj values, with lighter colors indicating greater statistical significance. The vertical red dashed line marks the significance threshold of P-Adj = 0.05. Terms in each cluster are ordered by enrichment significance.

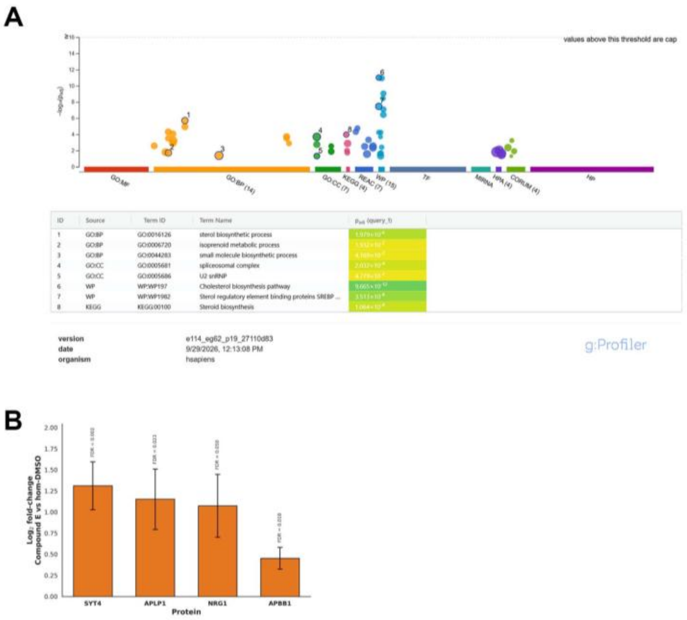


**Figure S5. Enrichment analysis and proteomic changes associated with Compound E.** **(A)** Functional enrichment analysis of proteins significantly altered by Atorvastatin relative to APP-hom DMSO, using g:Profiler. The analysis used all proteins detected in the experiment as the custom background. In the bubble plots, the y-axis shows enrichment significance (-log10 P-Adj value), and the x-axis lists significant terms grouped by data source. The table lists significant term identifications and P-Adj values. **(B)** $Log_2$ fold-changes for selected Compound E-associated proteins relative to APP-hom DMSO. Bars show mean $log_2$ fold-change, and error bars indicate 95% confidence intervals from Welch's t-tests. We calculated adjusted p-values using the Benjamini–Hochberg method across the Compound E proteome-wide comparison. Positive values indicate higher abundance following Compound E treatment. The bar plot is associated with results in Table 3.

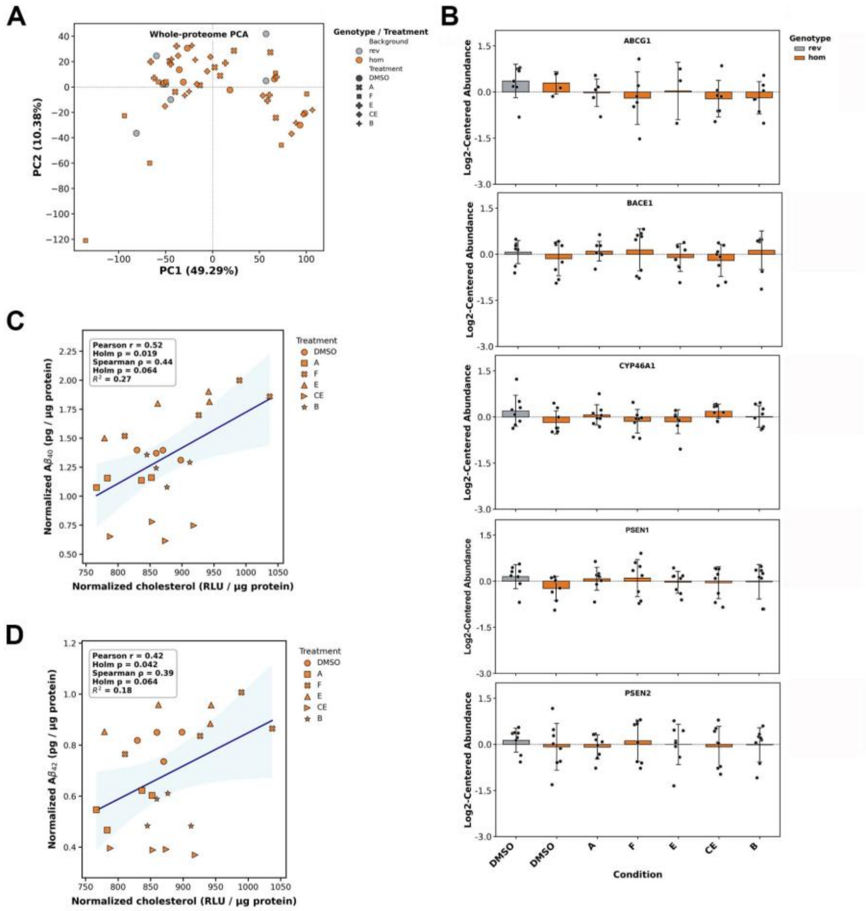


E

| Condition | Cholesterol (Mean ± SEM | Δ Chol vs DMSO (%) | Aβ40 (Mean ± SEM) | Δ Aβ40 vs DMSO (%) | Aβ42 (Mean ± SEM) | Δ Aβ42 vs DMSO (%) |
|---|---|---|---|---|---|---|
| DMSO | 864.31 ± 14.12 | 0.00% (Ref) | 1.37 ± 0.02 | 0.00% (Ref) | 0.81 ± 0.03 | 0.00% (Ref) |
| A | 809.49 ± 20.59 | -6.34% | 1.13 ± 0.02 | -17.33% | 0.56 ± 0.03 | -31.17% |
| F | 940.69 ± 48.99 | +8.84% | 1.77 ± 0.10 | +29.16% | 0.87 ± 0.05 | +6.72% |
| E | 881.25 ± 39.05 | +1.96% | 1.75 ± 0.09 | +28.09% | 0.91 ± 0.03 | +12.21% |
| CE | 858.34 ± 27.13 | -0.69% | 0.70 ± 0.04 | -48.99% | 0.39 ± 0.01 | -52.47% |
| B | 873.21 ± 14.44 | +1.03% | 1.24 ± 0.06 | -9.24% | 0.54 ± 0.03 | -33.41% |

**Figure S6. Pharmacological perturbation analysis to assess cellular cholesterol and amyloid-beta (Aβ) processing correlations. (A)** Principal component analysis (PCA) of pharmacological perturbation profiles. Proteomic variance along the first two principal components (PC1 vs. PC2) across cultured iNeurons treated with specific compounds or vehicle controls (listed at the end of this legend). Individual points represent independent culture conditions or replicates, illustrating global phenotypic clustering and separation driven by each condition**. (B)** Abundances of selected proteins presented as global log2-centered values (normalized across all retained acquisitions). Bars denote means with ± SD, and overlaid individual points represent independent biological replicates. The horizontal dashed line denotes the experiment-wide centered baseline (=0.0, corresponding to the mean abundance across all acquisitions). Correlation between intracellular cholesterol and Aβ-40 **(C)** and Aβ-42 secretion **(D).** Scatter plots evaluate relationships between normalized intracellular cholesterol and secreted Aβ species across APP-hom perturbation conditions. Individual points represent independent biological replicates, and distinct symbols indicate the compound tested. Solid lines denote ordinary least-squares linear regression trendlines, with shaded bands indicating the 95% confidence intervals of the mean fit. Insets report the Pearson correlation coefficient (r), coefficient of determination ($R^2$), Spearman rank correlation coefficient (p), and multiplicity-adjusted significance using the Holm–Bonferroni correction (Holm p). **(D)** Quantitative summary of RFU for cholesterol measurements, secreted Aβ40 and Aβ42 values across all evaluated pharmacological treatments relative to vehicle-treated control (APP-hom-DMSO). Values are reported as mean ± SEM across independent biological replicates (n = 4 per condition). Compound treatments include Atorvastatin (A), Fatostatin (F), Efavirenz (E), Compound E (CE), and β-Secretase Inhibitor IV (B).
